# Humans are primarily model-based learners in the two-stage task

**DOI:** 10.1101/682922

**Authors:** Carolina Feher da Silva, Todd A. Hare

## Abstract

Distinct model-free and model-based learning processes are thought to drive both typical and dysfunctional behaviours. Data from two-stage decision tasks have seemingly shown that human behaviour is driven by both processes operating in parallel. However, in this study, we show that more detailed task instructions lead participants to make primarily model-based choices that have little, if any, simple model-free influence. We also demonstrate that behaviour in the two-stage task may falsely appear to be driven by a combination of simple model-free and model-based learning if purely model-based agents form inaccurate models of the task because of misconceptions. Furthermore, we report evidence that many participants do misconceive the task in important ways. Overall, we argue that humans formulate a wide variety of learning models. Consequently, the simple dichotomy of model-free versus model-based learning is inadequate to explain behaviour in the two-stage task and connections between reward learning, habit formation, and compulsivity.

## Introduction

Investigating the interaction between habitual and goal-directed processes is essential to understand both normal and abnormal behaviour.^1–3^ Habits are thought to be learned via model-free learning,^4^ a strategy that operates by strengthening or weakening associations between stimuli and actions, depending on whether the action is followed by a reward or not.^5^ Conversely, another strategy known as model-based learning generates goal-directed behaviour,^4^ and may potentially protect against habit formation.^6^ Model-based behaviour selects actions by computing their current values based on a model of the environment.

Two-stage tasks have been used frequently to dissociate model-free and model-based influences on behaviour.^6–24^ Their critical feature is that participants make choices at the first stage of each trial, then transition probabilistically to a specific second-stage state (Figure 1A). Each of the two first-stage actions leads to one second-stage state with higher probability (e.g. 70%) and to the other with lower probability (e.g. 30%). Thus, there are common (high-probability) and rare (low-probability) transitions that depend on the first-stage choice. Model-free and model-based learning generate different predictions about the probability that in the next trial the participant will repeat their previous first-stage choice as a function of the previous transition and outcome. According to traditional logic, simple model-free agents are more likely to repeat a first-stage action that resulted in a reward, regardless of transition, and thus exhibit a positive main effect of reward (Figure 2A, although see^25^). Conversely, model-based agents first consider which second-stage stimulus is most likely to yield a reward, then select the first-stage action that will most likely lead to it, based on a model (i.e. knowledge) of the task’s structure.^7^ Thus, model-based agents factors in the transition and exhibit a positive reward by transition interaction effect (Figure 2B). Hybrid agents that combine both strategies exhibit both effects (Figure 2C).

**Figure 1:**
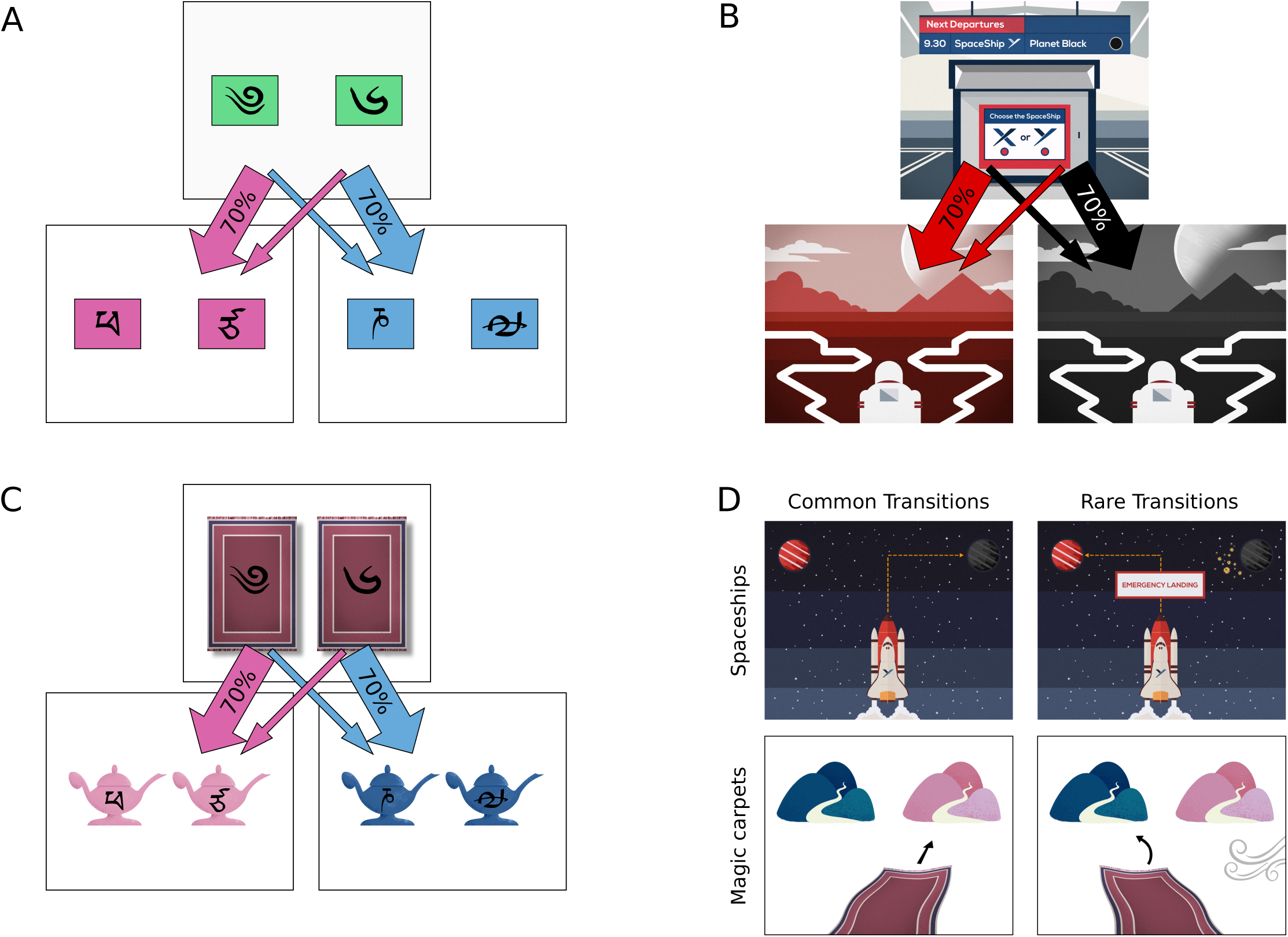
The stimuli used in the three versions of the two-stage task. A: The original, abstract version of the task used by Daw et al.^7^ In each trial, the participant makes choices in two consecutive stages. In the first stage, the participant chooses one of two green boxes, each of which contains a Tibetan character that identifies it. Depending on the chosen box, the participant transitions with different probabilities to a second-stage state, either the pink or the blue state. One green box takes the participant to the pink state with 0.7 probability and to the blue state with 0.3 probability, while the other takes the participant to the blue state with 0.7 probability and to the pink state with 0.3 probability. At the second stage, the participant chooses again between two boxes containing identifying Tibetan characters, which may be pink or blue depending on which state they are in. The participant then receives a reward or not. Each pink or blue box has a different reward probability, which randomly changes during the course of the experiment. The reward and transition properties remain the same in the versions of the two-stage task shown in B and C. B: Spaceship version, which explains the task to participants with a story about a space explorer flying on spaceships and searching for crystals on alien planets. C: Magic carpet version, which explains the task to participants with a story about a musician flying on magic carpets and playing the flute to genies, who live on mountains, inside magic lamps. D: Depiction of common and rare transitions by the magic carpet and spaceship tasks. In the magic carpet task training session, common transitions are represented by the magic carpet flying directly to a mountain, and rare transitions are represented by the magic carpet being blown by the wind toward the opposite mountain. In the spaceship task, common transitions are represented by the spaceship flying directly to a planet, and rare transitions are represented by the spaceship’s path being blocked by an asteroid cloud, which forces the spaceship to land on the other planet.

**Figure 2:**
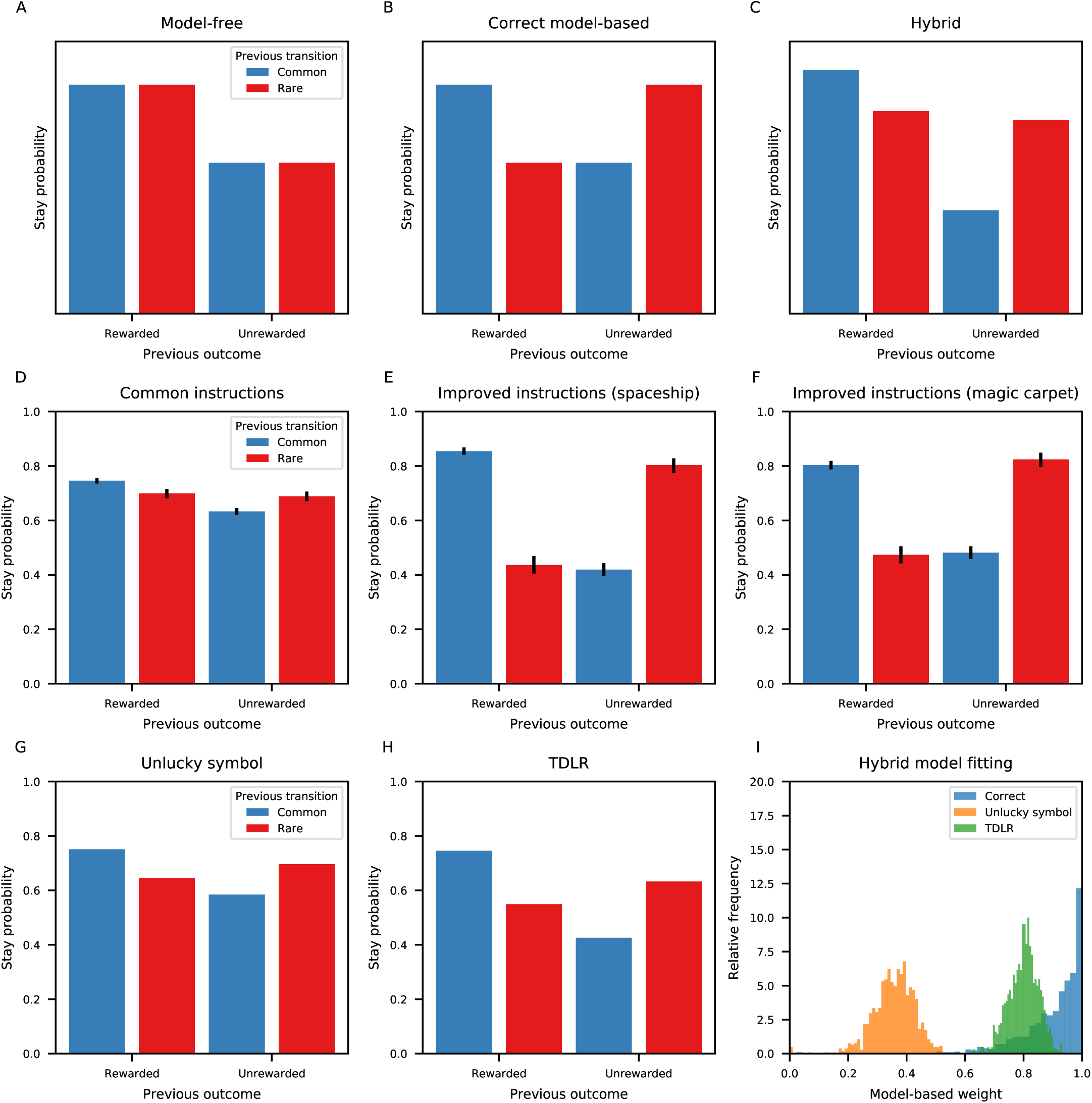
Stay probabilities for human participants; stay probabilities and model-based weights for simulated agents. Top row: Idealized stay probabilities of purely model-free (A), purely model-based (B), and hybrid (C) agents as a function of the previous outcome and transition. To generate the hybrid data plotted in this figure, we used the logistic regression coefficient values for data from adult participants in a previous study.^20^ Middle row: D) The behaviour of participants (*N* = 206) after receiving common instructions^21^ shows both a main effect of reward and a reward by transition interaction. In contrast, the choices of participants after receiving improved instructions in the new E) spaceship (*N* = 21) and F) magic carpet (*N* = 24) tasks show a much more model-based pattern. Bottom row: Purely model-based agents that use incorrect models of the two-stage task can look like hybrid agents that use a combination of model-based and model-free learning. Panels G and H show the mean stay probabilities for each type of model-based agent. I) The histograms show the fitted model-based weight parameters (*w*) for simulated agents using the correct (blue; median = 0.94, 95% interval = [0.74, 1.00]), unlucky-symbol (orange; median = 0.36, 95% interval = [0.24, 0.48]), and transition-dependent learning rates (TDLR) (green; median = 0.80, 95% interval = [0.70, 0.90]) models of the task. We simulated 1 000 agents of each type. Model-based weights for each agent were estimated by fitting the simulated choice data (1 000 choices per agent) with the original hybrid model by maximum likelihood estimation. Error bars in panels D to F represent the 95% highest density intervals. Error bars for the simulated data are not shown because they are very small due to the large number of data points.

Past studies employing the original two-stage task (Figure 1A) have always reported that healthy adult humans use a hybrid mixture of model-free and model-based learning (e.g.^7, 20, 21^ and Figure 2D). Moreover, most studies implementing modifications to the two-stage task designed to promote model-based learning^21, 22, 24^ find a reduced but substantial model-free influence on behaviour. Overall, the consensus has been that the influence of simple model-free learning on human behaviour is ubiquitous and robust.

Our current findings call into question how ubiquitous simple model-free learning is. We found that slight changes to the task instructions markedly reduced the apparent evidence for model-free learning in two separate experiments (Figure 2E–F). We ran a series of simulations of purely model-based agents that used incorrect models of the two-stage task, and found that the agents falsely appeared to be hybrid. Lastly, we show that human participants often misrepresent basic features of the two-stage task. This means inaccurate models of the two-stage task could be the true reason for some, or even all, of the past findings of hybrid behaviour rather than competition between model-free and model-based learning algorithms.

## Results

### Improving the two-stage task instructions decreases the apparent model-free influence on behaviour

We developed two modified versions of the two-stage task—the magic carpet task and the spaceship task—with the goal of clearly explaining all features of the task. We thought that if participants’ apparent model-free behaviour was caused by a poor mental representation of the task, our improved instructions would shift them toward the correct model-based behaviour. Conversely, if their apparent model-free behaviour truly resulted from a competition between parallel model-based and model-free systems, our improved instructions would not make any difference.

Specifically, we incorporated a detailed story in the instructions and stimuli (Figure 1B–D). In the magic carpet task, the participant chose a magic carpet and flew on it to Pink or Blue Mountain, where genies might give them a gold coin. In the spaceship task, the participant bought a spaceship ticket and flew to Planet Red or Black in search of valuable crystals that grew inside obelisks. Previous studies have already used stories to explain the two-stage task to human participants,^20, 21^ but they did not provide a reason for all the events in the task (see Supplementary Methods). Conversely, our instructions provided a concrete reason for every potential event within the task; in particular, different explanations and visual depictions were given for common versus rare transitions (Figure 1D) during training. Importantly, the magic carpet task used different stimuli for the training trials versus the main task, and there were no visual depictions of the transitions during the main task. During the practice trials, we also displayed additional messages, which explained the reason for each event again as it happened and could not be skipped (see Methods).

Both the magic carpet and the spaceship tasks have the same payoff structure and transition probabilities as the original two-stage task.^7^ The magic carpet task replicates all features of the original two-stage task, except for the instructions. In contrast, the spaceship task was originally designed to test how humans learn when the first-stage states are defined by a combination of two stimuli. In this task, there were four different first-stage stimuli (“flight announcements”) that defined the initial state and the transition probabilities associated with the two first-stage actions (Figure 4A). This feature allowed us to demonstrate that a common reverse inference—significant main effects of reward in a logistic regression analysis indicate model-free learning—is invalid.

**Figure 3:**
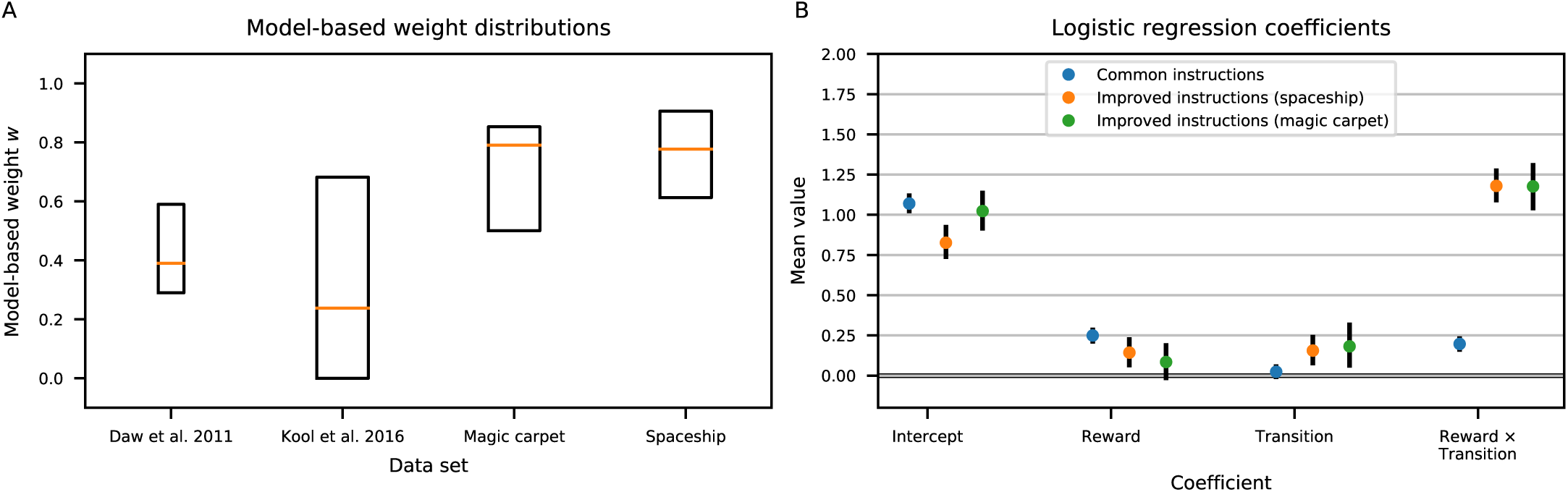
Model-based weights and logistic regression coefficients for different empirical data sets. A) Summary statistics (25%, 50%, and 75% percentiles) of the estimated model-based weights for four data sets: Daw et al.^7^ (*N* = 17), Kool et al.^21^ (common instructions data set, *N* = 206), the spaceship data set (*N* = 21), and the magic carpet data set (*N* = 24). For the Daw et al.^7^ data set, the weight estimates were simply copied from the original article. For the other data sets, we obtained the weight estimates by a maximum likelihood fit of the hybrid reinforcement learning model to the data from each participant. B) This plot shows the mean and 95% highest density intervals of all coefficients in the hierarchical logistic regressions on stay probabilities for the common instructions,^21^ magic carpet, and spaceship data sets. These logistic regression coefficients were used to calculate the stay probabilities shown in Figure 2D–F. The improved instruction data sets have both lower Reward and higher Reward x Transition interaction effects compared to the common instructions data set. Note that the main effect of reward in the spaceship task is actually inconsistent with a model-free influence on behaviour in that task (see Figure 4). There are significant effects of transition on stay probabilities in the improved instruction data sets as well. This coefficient indicates that the probability of repeating the same first-stage action increases after a common transition and decreases after a rare transition to the second stage. Transition effects are consistent with standard model-free, model-based, or hybrid behaviour under certain combinations of the learning and choice parameters in those algorithms (see Supplementary Results).

**Figure 4:**
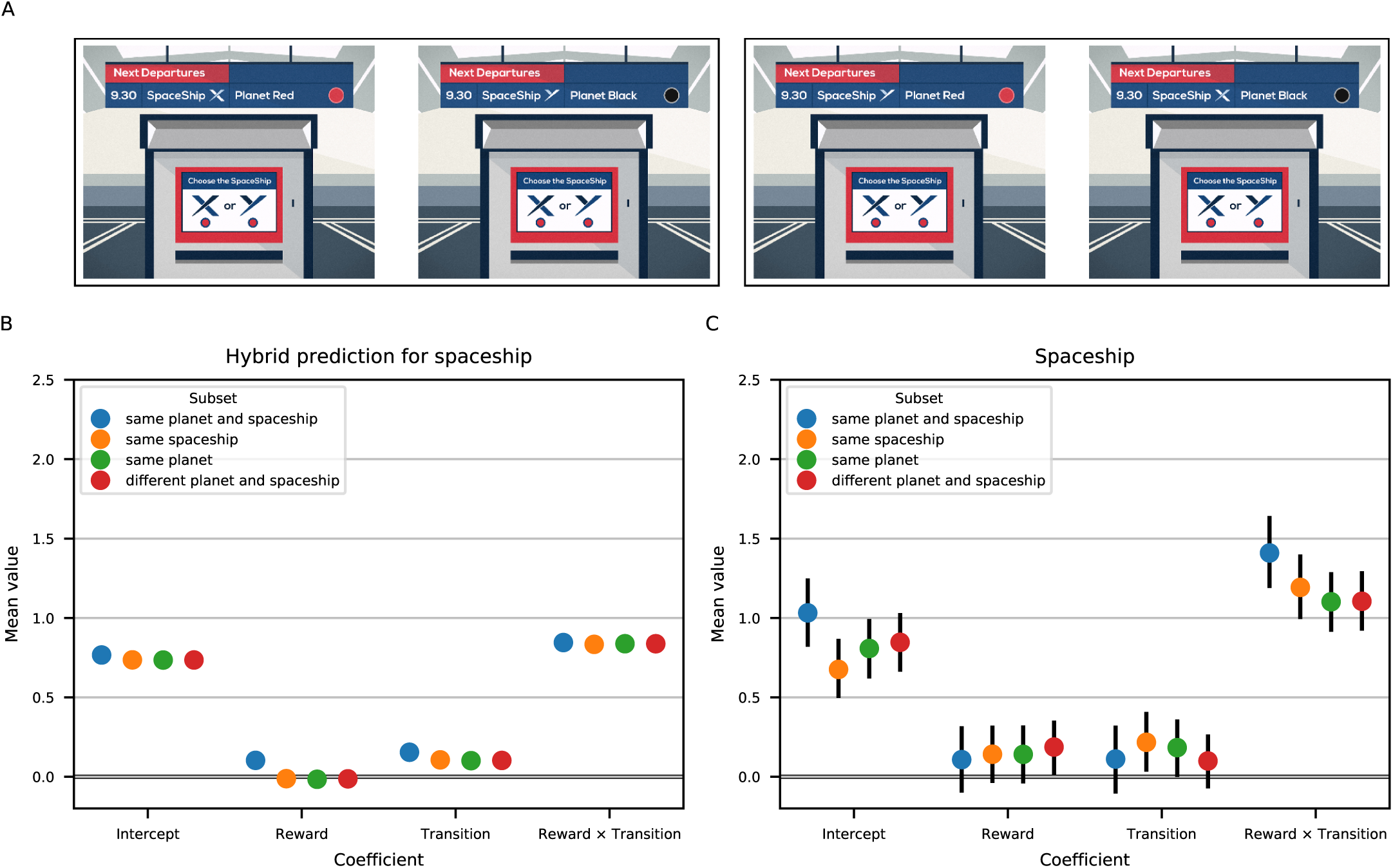
Example of a reward main effect that cannot be driven by model-free learning. A) Each trial of the spaceship task began with a flight announcement on an information board above a ticket machine. The name of a spaceship was presented for 2 seconds, then the name of the planet the spaceship was scheduled to fly to appeared for another 2 seconds. There were two spaceships (X and Y) and two planets (Red and Black), and hence four possible flight announcements. The participant selected a spaceship to fly on based on the announced flight. Each announcement listed only one flight, but still provided complete information about all flights. This is because the spaceships always departed at the same time and flew to different planets: If one would fly to Planet Red, then the other would fly to Planet Black and vice versa. Thus, the two screens in each rectangle of panel A are equivalent. B) Results for simulated hybrid agents (*N* = 5000) performing 1 000 trials of the spaceship task under the standard assumption that model-free learning does not generalize between different state representations (i.e. different flight announcements). Simulation parameters were the median estimates obtained by fitting the hybrid model to the human spaceship data by maximum likelihood estimation. The points on the plot represent coefficients from a logistic regression analysis on the simulated choices, with consecutive trial pairs divided into four categories: (1, blue) same planet and spaceship, (2, orange) same spaceship, (3, green) same planet, and (4, red) different planet and spaceship. This division was made with regard to which flight was announced in the current trial compared to the preceding trial. Error bars are not shown because they are very small due to the large number of data points. C) Logistic regression results for the human spaceship data (*N* = 21), with consecutive trial pairs divided into the same four categories. In contrast to the simulated hybrid agents, the small reward effect in human behaviour does not differ across categories. Thus, it is inconsistent with a standard model-free learning system. Error bars represent the 95% highest density intervals.

#### Hybrid model fits indicate that behaviour becomes more model-based with comprehensive instructions

To test the impact of our instructions on the apparent levels of model-based and model-free behaviour, we fitted to the data the standard hybrid reinforcement learning model, proposed originally by Daw et al.^7^ In order to facilitate comparison with previous studies, we fit the model to each participant using maximum likelihood estimation. The hybrid model combines the model-free SARSA(*λ*) algorithm with model-based learning and explains first-stage choices as a combination of the model-free and model-based state-dependent action values, weighted by a model-based weight *w* (0 *≤ w ≤* 1). A model-based weight equal to 1 indicates a purely model-based strategy and, conversely, a model-based weight equal to 0 indicates a purely model-free strategy.

The estimated model-based weights for the participants who performed the spaceship or the magic carpet task were substantially higher than the estimated weights obtained in two previous studies^7, 21^ using common (less comprehensive) task instructions (Figure 3A). We also fit a Bayesian hierarchical model containing the hybrid algorithm to the data from our magic carpet and spaceship tasks as well as the control condition data from a previous study using common instructions (*N* = 206).^21^ Our results indicate that the posterior probability that the average weights in the magic carpet and spaceship data sets are greater than the average weight in the common instructions data set is greater than 0.9999.

#### Standard logistic regression analyses also indicate primarily model-based behaviour in the magic carpet and spaceship tasks

We also analysed our results using a logistic regression analysis of consecutive trial pairs. In this analysis, the stay probability (probability of repeating a first-stage action) is a function of two variables: reward, indicating whether the previous trial was rewarded, and transition, indicating whether the previous trial’s transition was common or rare. Model-free learning generates a main effect of reward (Figure 2A), while model-based learning generates a reward by transition interaction (Figure 2B). The core finding in most studies is that healthy adult participants behave like hybrid agents (Figure 2C), exhibiting both a main effect of reward and a reward by transition interaction.

For comparison to our data, we re-analysed the common instructions data set (control condition from ^21^). Figure 3B shows that when all trial pairs were combined, the coefficient of the reward by transition interaction that indicates correct model-based control is 5.9 times larger in the magic carpet (95% HDI [4.5, 7.3]) and spaceship (95% HDI [4.7, 7.2]) tasks compared to the common instructions data.^21^ Moreover, the reward effect, which is generally considered an evidence of model-free control, is higher in the common instructions results (95% HDI [0.20, 0.30]) compared to the magic carpet results (95% HDI [− 0.03, 0.20]) with 0.99 probability (Bayes Factor: 145) and the spaceship results (95% HDI [0.05, 0.24]) with 0.98 probability (Bayes Factor: 51).

#### Spaceship task data reveal misleading evidence of model-free influence

Although choices in spaceship tasks showed large reward by transition interaction effects, there was a small, but significant main effect of reward on choices (Figure 3B). This may at first suggest that our enhanced instructions decreased but did not eliminate the influence of model-free learning on these participants. We took advantage of specific properties of the spaceship task to further investigate this reward effect. We found that it is misleading as evidence of model-free influence, because it contradicts one of the basic properties of model-free learning.

Within the spaceship task there are pairs of first-stage stimuli that indicate the same initial state by presenting different information (Figure 4A). This allows us to subdivide consecutive trial pairs into four categories, based on the information announced on the flight board. This information determines which first-stage action will most probably lead to a given second-stage state (Figure 4A and Table 1). We analysed the data from each category using the standard logistic regression.

**Table 1:** Reward main effects in different trial pair categories for the spaceship task. Pairs of consecutive trials were divided into four categories depending on the flight announcement, and each subset was separately analysed by logistic regression. The categories are numbered in the order (left to right) that they are shown in Figure 4. Column two lists the group-level means for the reward main effect by category. The values in square brackets denote the range of the 95% highest density interval (HDI). Column three lists the posterior probability that the main effect of reward is greater than zero in each category and its equivalent Bayes Factor. Column four lists the (dis)similarities in the flight announcement screens across consecutive trial pairs that are used to define categories 1–4. For categories 2 and 3 combined, which corresponded to “stay” choices requiring the selection of a different spaceship, the posterior probability that the reward effect is greater than zero is 98% (Bayes Factor: 46.0).

| Category | Reward effect magnitude | Probability $> 0$<br>(Bayes Factor) | Announcements included |
| --- | --- | --- | --- |
| 1 | 0.12 $[-0.10, 0.32]$ | 85% (5.6) | The same spaceship and planet |
| 2 | 0.14 $[-0.03, 0.32]$ | 94% (14.9) | The same spaceship, but different planets |
| 3 | 0.14 $[-0.04, 0.33]$ | 93% (13.9) | Different spaceships, but the same planet |
| 4 | 0.19 $[+0.02, 0.36]$ | 98% (62.9) | Different spaceships and different planets |

If the reward effect was driven by model free learning, it should be positive only when the stimulus-response pairing is identical across trials (category 1, same spaceship and planet). This is because model-free learning is assumed to be unable to generalize between distinct state representations^15, 16, 21^ and thus should have no influence on choices in categories 2–4 (Figure 4B). The results, however, are contrary to this expectation (Figure 4C): The observed reward effect had a similar magnitude in all four categories, including those where either the stimuli or the response required to implement the same choice are different between two consecutive trials. Note that generalization between different flight announcements implies knowledge about the task structure, because the flight announcements define the transitions between first- and second-stage states. These results strongly suggest that the observed reward effects are not model-free. Instead, they are more likely model-based, except that the models participants used were not completely correct, as assumed by the analysis.

### The logistic regression model is better than the hybrid model at explaining first-stage choices in all tasks

We found that the hybrid reinforcement learning model^7^ does not describe humans’ first-stage choices well in any of the three data sets. We compared how well three models—the simple logistic regression model of stay probabilities in consecutive trial pairs, the hybrid model, and the correct model-based model—fit human participants’ first-stage choices. We compared the models based on how well they explained first-stage choices only, because the logistic regression model is only fit to first-stage choices, and the model-free and model-based algorithms only generate different behaviour at the first stage. Both AIC and PSIS-LOO metrics strongly indicated that the simple logistic regression model best explained participants choices (Table 2). In fact, PSIS-LOO scores favoured the logistic regression model by more than four standard errors.

**Table 2:** Overall AIC and PSIS-LOO scores for three models fitted to human participant data. Participants belonged to either the common instructions (*N* = 206), magic carpet (*N* = 24), or spaceship (*N* = 21) condition. We fit participants’ choice data using a logistic regression model of consecutive trial pairs, the correct model-based reinforcement learning model, and the hybrid reinforcement learning model. AIC calculation failed for one participant in the common instructions condition and so this participant was excluded from the AIC analysis. The difference between PSIS-LOO scores for the logistic regression and the hybrid model were −565.0 *±* 104.6, −256 *±* 60.2, and −507.2 *±* 63 for the common instructions, magic carpet, and spaceship data sets respectively. Errors given for the PSIS-LOO scores correspond to the standard error. For both the AIC and PSIS-LOO metrics, smaller values indicate better model fits.

| Condition | AIC |  |  | PSIS-LOO |  |  |
| --- | --- | --- | --- | --- | --- | --- |
|  | Logistic regression | Model-based | Hybrid | Logistic regression | Model-based | Hybrid |
| Common instructions | 27073 | 27441 | 27246 | 26725.7 ± 140.4 | 27685.0 ± 152.8 | 27290.8 ± 158.6 |
| Magic carpet | 4682 | 5113 | 5063 | 4665.7 ± 67.8 | 5032.5 ± 63.2 | 4921.8 ± 63.5 |
| Spaceship | 5118 | 5684 | 5849 | 5109.9 ± 73.3 | 6415.0 ± 85.7 | 5617.2 ± 66.0 |

The logistic regression model describes stay probabilities as constant throughout the experiment; it is not a learning model. This suggests that participants’ behaviour in the two-stage task is not well described as a combination of the correct model-based and model-free learning algorithms that constitute the hybrid model. Notably, the logistic regression model revealed little evidence of simple model-free learning with our improved instructions. Overall, our results suggest that participants in the magic carpet and spaceship tasks used a slightly incorrect model-based strategy. (See also Supplementary Results.)

### Model-based learning can be confused with model-free learning

Our experiments with improved instructions demonstrate that participants’ understanding of the task plays a large role in how model-based or model-free they appear to be. Here, we present simulated data to demonstrate that purely model-based agents using incorrect task models may be misclassified as hybrid when the data are analysed by either of the standard methods: logistic regression or reinforcement learning model fitting. We present two examples of incorrect task models. We do not suggest that they are the only or even the most probable ways that people may misconceive of the task. Instead, they are merely examples to demonstrate our points.

As a reference, we simulated correct model-based agents, based on the hybrid model proposed by Daw et al. with *w* = 1.^7^ Consistent with recent work by Sharar et al.,^26^ even when agents have a *w* equal to exactly 1, used the correct model of the task, and performed 1 000 trials, the recovered *w* parameters were not always precisely 1 (Figure 2I). This is expected, because parameter recovery is noisy and in the standard specification of the hybrid model *w* cannot be greater than 1, thus any error was an underestimate of *w*.

The two alternative, purely model-based learning algorithms are: the “unlucky symbol” and the “transition-dependent learning rates” (TDLR) algorithm (see Methods for full details). Briefly, the unlucky symbol algorithm has the mistaken belief that certain first-stage symbols decrease the reward probability of second-stage choices. In the current example, we simulated agents that believe a certain first-stage symbol is unlucky and lowers the values of second-stage actions by 50%. The TDLR algorithm is a model-based learning algorithm that has a higher learning rate after common than rare transitions. Note that both algorithms contain the correct state transition structure. This suggests that checking if participants understood the transitions is not enough to show that they properly modeled the task; there may be other elements of their models that are incorrect.

Our simulations show that TDLR and unlucky symbol agents, despite being purely model-based, display the same behavioural pattern as healthy adults given the common task instructions and simulated hybrid agents (Figure 2). We analysed the simulated data with the hybrid algorithm containing the correct model of the task (i.e. the standard analysis). The resulting distributions of the estimated model-based weights are shown in Figure 2I. They indicate a model-free influence on the behaviour of purely model-based TDLR and unlucky-symbol agents. Moreover, when fitting the correct model-based and hybrid models to the data and performing model comparisons, we found lower AIC scores (indicating a better fit; difference = −5 897) for the purely model-based algorithm only when agents used the correct model. For the TDLR and unlucky-symbol agents, the correct model-based algorithm yielded higher AIC scores (i.e. a worse fit) than the hybrid model (AIC differences = 26 858 and 198 932, respectively) even though these agents did not include any model-free influence. Together, these results demonstrate that analyzing two-stage task choices using a hybrid algorithm can lead to the misclassification of purely model-based agents as hybrid if the agents have an incorrect model of the task.

### Human behaviour deviates from the hybrid model’s assumptions

We tested if human behaviour following common instructions^21^ also violates assumptions of the hybrid model. First, we note that poor overall hybrid model fits were significantly associated with more apparent evidence of model-free behaviour. There was a significant positive correlation between the model-based weight and the log-likelihood of the model fit (Spearman’s rho = 0.19, *P* = 0.005). There is no inherent reason for such correlation if participants are using a hybrid mixture of simple model-free and correct model-based learning, assuming all other factors (e.g. exploration) are equal. Upon further analysis, we found that this correlation was driven by participants with an estimated model-based weight smaller than 0.1 (about 43% of the sample). This suggests that human participants that behaved in a way that substantially deviated from the hybrid model’s assumptions were assigned very low model-based weights in order to maximize the correspondence between hybrid model predictions and their behaviour (although this was still a poor match). One potential reason for this bias is that the model-free portion of the hybrid model has two unique free parameters, giving it flexibility to match different behaviours, while the model-based portion has none. Therefore, the model-free portion of the hybrid algorithm may be better at fitting behaviour that deviates from the correct model.

Second, the standard hybrid model cannot explain why participants in this experiment behaved differently when first-stage symbols were presented on different sides of the screen in consecutive trials. In many implementations of the two-stage task, the symbols presented at each stage appear on random sides. The sides are irrelevant and only the symbol identity matters. If participants understand the task, changes in first-stage symbols locations should not influence their choices. Nevertheless, past studies have anticipated that participants might have a tendency to repeat key presses at the first stage regardless of symbol locations and thus modified the standard hybrid model to add a response stickiness parameter.^21^ For comparison with participant data, we simulated hybrid agents with response stickiness. We then divided consecutive trial pairs from the simulated and participant data sets into two subsets: (1) same sides, if the first-stage choices were presented on the same sides in both trials, and (2) different sides, if the first-stage choices switched sides from one trial to the next. We analysed each subset separately using logistic regressions (Figure 5A and C). The results show a larger intercept in the same-sides subset compared to the different-sides subset for both simulated agents and participants. In the simulated data, this effect was caused by response stickiness. However, human participants also showed a larger reward coefficient in the same-sides subset (mean 0.41, 95% highest density interval (HDI) [0.35, 0.47]) versus the different-sides subset (mean 0.14, 95% HDI [0.08, 0.19]), with the posterior probability that the reward coefficient is larger in the former being greater than 0.9999 (Bayes Factor: *>* 60 000).

**Figure 5:**
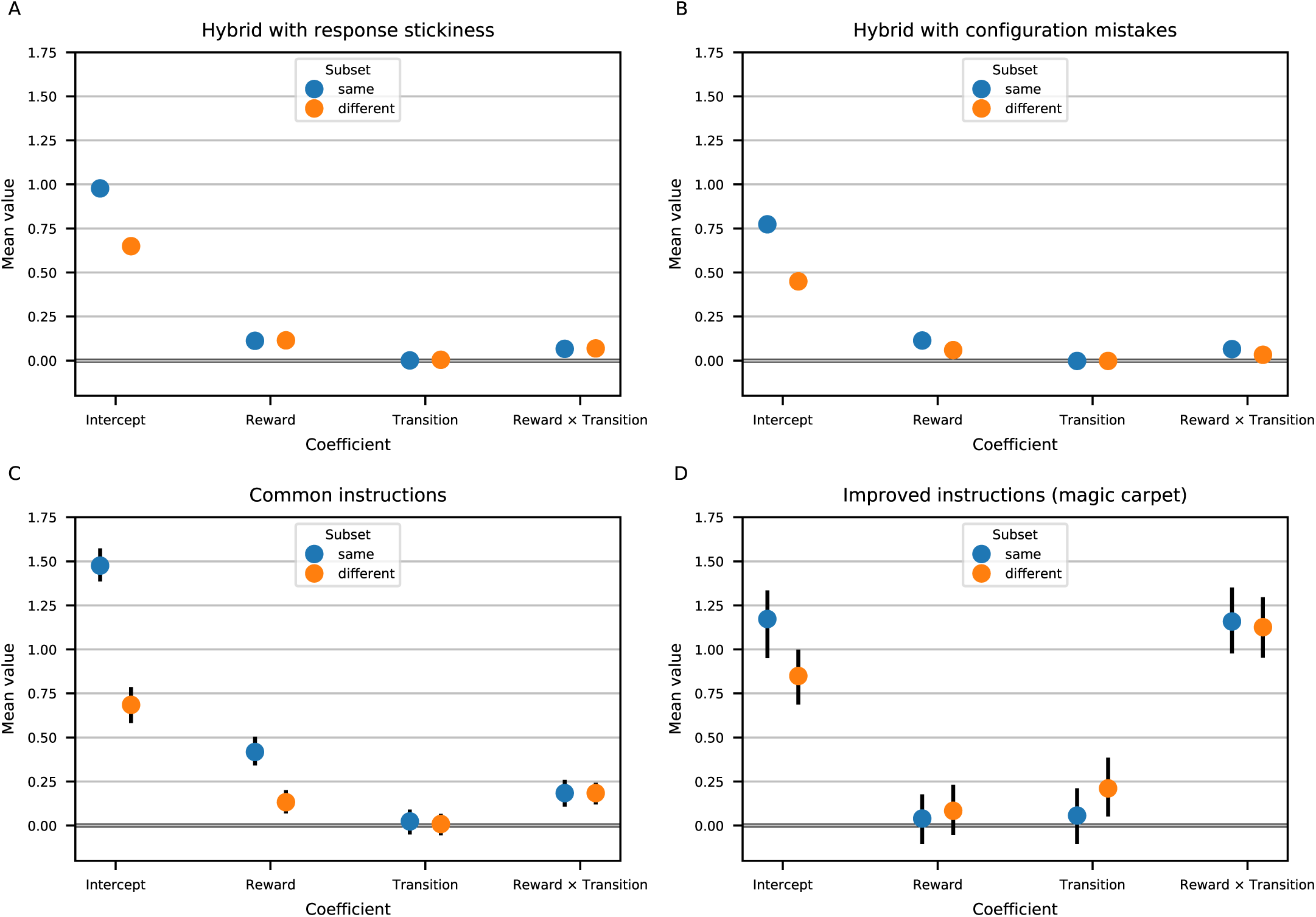
Simulated agents’ and real participants’ behaviour can be influenced by irrelevant changes in stimulus position. All four panels show behaviour in the two-stage task on trials in which the position of the first-stage stimuli remains the same (blue) across consecutive trials compared to trials in which the first-stage stimuli are in different (orange) positions from one trial to the next. The coefficients in these graphs are from a logistic regression analysis explaining stay probabilities on the current trial as a function of reward, transition, and the interaction between reward and transition on the previous trial. In contrast to previous studies, we analysed the data after dividing the trials into two categories based on whether or not the positions of the first-stage stimuli were the same or different across consecutive trials. A) Results from a simulation of hybrid agents with response stickiness (*N* = 5 000) performing 1 000 trials of the two-stage task. The median parameter values from the common instructions data set^21^ were used in these simulations. B) Results from a simulation of hybrid agents that occasionally made configuration mistakes (*N* = 5 000) performing 1 000 trials of the two-stage task. The median parameter values from the common instructions data set^21^ and a 20% configuration mistake probability were used in these simulations. Error bars are not shown for these simulated results because they are very small due to the large number of data points. C) Results from a re-analysis of the common instructions data set^21^ (*N* = 206). This study used story-like instructions, but did not explicitly explain why the stimuli might be on different sides of the screen from trial to trial. The Reward effect significantly changed between trial-type subsets in this data set. D) Results from the magic carpet task (*N* = 24), which provided explicit information about why stimuli may change positions across trials and that these changes were irrelevant for rewards and transitions within the task. There were no significant differences in the regression coefficients between the two subsets of trials on this task. Error bars in panels C and D represent the 95% highest density intervals.

There are several potential explanations for these side-specific results. It could be that model-free learning is sensitive to stimulus locations or specific responses and considers that each symbol has a different value depending on where it is presented or on which key the participant has to press to select it. In this case, the reward effect for the different-sides subset should be zero, but it is not. Another possibility is that when the sides switched, participants were more likely to make a mistake and press the wrong key, based on the previous rather than current trial configuration. To further investigate this latter possibility, we fit these data with a hybrid model that included an extra parameter quantifying the probability of making a configuration mistake and pressing the wrong key after a location change (see Methods subsection “Fitting of reinforcement learning models” for details). Note that this configuration mistake parameter is distinct from decision noise or randomness, because it quantifies response probabilities when symbols have switched places from one trial to the next and thus only decreases effect sizes in the different-sides subset (Figure 5B).

When looking at individual participants, distinct types of performance were readily apparent. Out of 206 participants, 111 had an estimated probability of making a configuration mistake lower than 0.1 (based on the median of their posterior distributions). In contrast, 51 participants had an estimated configuration-mistake probability higher than 0.9. The goodness of fits for the two models were equivalent on average for the 111 low-configuration-mistake participants (mean score difference: 0.0, standard error: 0.1), As expected, the configuration mistake model fit better for the 51 high-configuration-mistake participants (mean score difference: −34.1, standard error: 6.2). These results suggest that most participants rarely made configuration mistakes, but approximately 25% made mistakes more than 9 out of 10 times when the symbols switched sides.

However, it is possible that the high-configuration-mistake participants were not truly making mistakes. Configuration mistakes decrease all effect sizes in the different sides subset, including the reward by transition interaction effect (Figure 5B), but this was not observed in the common instructions data set. Rather, some participants may have instead mismodeled the task—for example, they may have believed a stimulus’s location was more important than its identity. In any case, these results suggest that different participants conceive of the two-stage task in different ways, and that many participants misconceptualized basic aspects of the task.

#### Smaller side-specific effects and fewer configuration mistakes with enhanced magic carpet instructions

In contrast to the common instructions sample,^21^ participants who performed the magic carpet task showed little difference in the logistic regression coefficients between the same-side and different-side subsets (Fig. 5D). They also made few configuration mistakes, with only one participant having a mistake probability greater than 0.05. Thus, the enhanced magic carpet instructions vastly increased the probability that participants would act correctly when equivalent choices were presented on different sides of the screen.

## Discussion

We show that simple changes in the two-stage task instructions led healthy adult humans to behave in a highly model-based manner. This is in contrast to most studies, which suggest that decisions in these tasks are driven by a combination of model-based and model-free learning. However, we also show that if purely model-based agents use incorrect models, analysis results can falsely indicate an influence of model-free learning and mistakenly classify them as hybrid. If participants use incorrect models of the task, none of the traditional markers of model-based or model-free behaviour derived from logistic regression or hybrid algorithms indicate the ground-truth about participants’ strategies. Therefore, our work, together with other recent reports on the limitations of hybrid model fits,^26, 27^ indicates the need to reconsider aspects of both the empirical tools and theoretical assumptions that currently pervade the study of reward learning.

Human behaviour does not necessarily fall into the dichotomy of simple model-free and correct model-based learning assumed in the design and analysis of two-stage tasks.^28–35^ Instead, agents can act on a multitude of strategies (Figure 6). It is known that this multidimensional space includes complex model-free algorithms that can mimic model-based actions in some cases.^29, 31^ We show that it also includes model-based strategies that can appear hybrid because they are inconsistent with the environment. This adds uncertainty to any attempt to classify strategies in the two-stage task.

**Figure 6:**
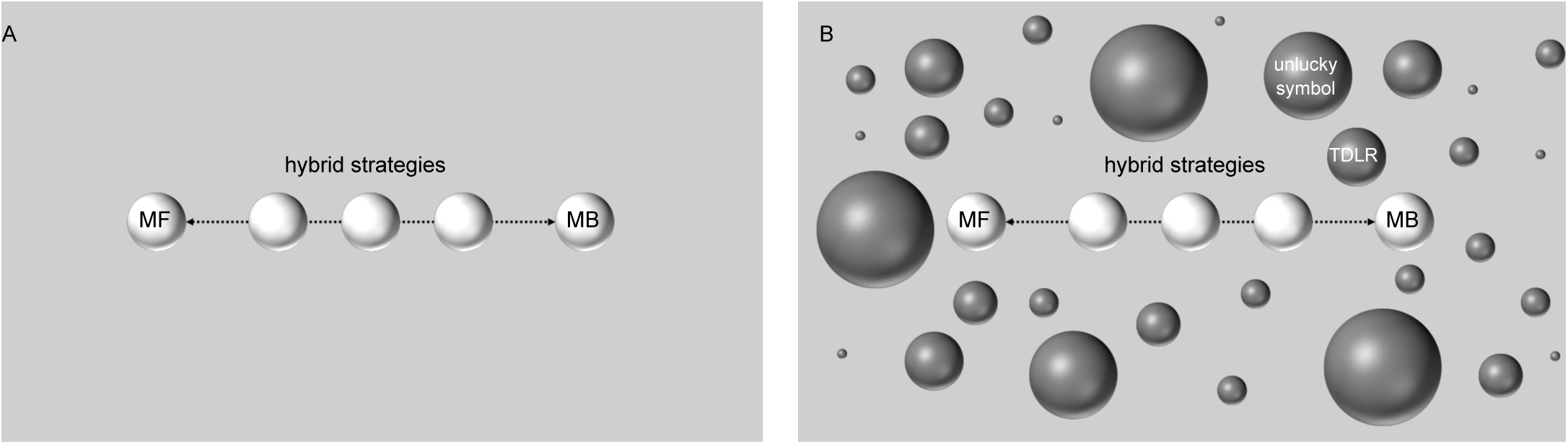
Simplified diagrams representing the strategy space in the two-stage task. A) The commonly employed analysis methods for two-stage task behaviour assume that the space of all strategies agents can employ to perform this task contains only the model-free (MF) and correct model-based (MB) strategies, as well as intermediate hybrid strategies on the line between them. B) However, the true strategy space also contains other purely model-based strategies, such as the transition-dependent learning rates (TDLR) and unlucky symbol strategies, among countless others. If these other strategies are projected onto the line segment between model-free and correct model-based behaviour, they will yield the false conclusion that participants are using a hybrid MF/MB strategy. This is essentially what the standard analyses of the two-stage task do. In reality, a human participant who misconceives of the two-stage task in some way may use any of the potential strategies that exist in a multidimensional space. Importantly, if people misconceive the task in different ways, then they will use different incorrect models that correspond to their current mental representation of the task. Moreover, people may well switch between different strategies over the course of the experiment if they determine that their conceptualization of the task was wrong in some way. Unless we can ensure a complete understanding of the correct model-based strategy a priori, the vast space of possible incorrect models, heterogeneity across participants, and potential for participants to change models over time make accurate identification of a hybrid mixture of model-free and model-based learning a daunting, if not impossible, task.

Concluding that behaviour is a mix of two exemplar algorithms is especially tenuous. The observed choices do not match the predictions of either algorithm; arguably the most plausible reason for this is that participants are not using either algorithm. Still, it is possible that both algorithms operate in parallel and jointly influence decisions. Indeed, this has been the usual conclusion, probably because of strong prior beliefs in the existence of dual learning systems. However, it relies on the strong assumption that participants have an accurate task model, which we show may not hold in many cases.

In line with the possibility that apparently hybrid behaviour is, instead, driven by misconceptions of the task, we found drastic shifts toward (correct) model-based behaviour in healthy adult humans when participants received more elaborate instructions. It comes as no surprise that people don’t always understand or follow instructions well. However, the impact of these misconceptions on our ability to make accurate inferences about reward learning was unexpected to us. In both our local samples and openly available data, we found strong indications that people misconstrued the two-stage task. Among them are the location effects detected in the common instructions data.^21^ Those participants completed the task online, but similar behaviour has been reported by Shahar et al. from participants in a laboratory.^36^ Shahar et al. proposed that location effects are due to model-free credit assignment to outcome-irrelevant task features, such as stimulus locations and key responses.^36^ However, improved instructions for our tasks greatly reduced side-switch errors, suggesting that model-based misconceptions were more likely the cause. This result also argues against the alternative explanation that our instructions, rather than alleviating confusion, simply encouraged participants to be more model-based. While they might do so by explicitly describing and providing reasons for the transitions, the near elimination of location effects is direct evidence that they also facilitated the formation and use of more accurate task models.

Our spaceship task provides empirical evidence that healthy adult humans may employ partially incorrect models—even with improved instructions. Behaviour on that task was almost correct model-based. However, there was a small reward effect in a logistic regression analysis of consecutive trials with the same or different initial states. Such an effect is assumed to be evidence of model-free learning,^7^ but model-free learning is thought to be unable to generalize between distinct state representations.^15, 16, 21, 29^ If it drove the small reward effect in the spaceship task, it would be able to generalize not only between distinct state representations but also distinct actions. Therefore, the observed effect was more likely generated by imprecise model-based learning. It is not clear how the participants misunderstood the spaceship task, or even if they all misunderstood it in the same way. The fact that there are unlimited incorrect model-based models and some seem to mimic a hybrid agent is the core obstacle to drawing accurate conclusions from behaviour in the two-stage task. We also found that when the hybrid model doesn’t explain human choices well, it is biased toward indicating more model-free influence. These findings corroborate previous reports that a measure of understanding, the number of attempts required to pass a comprehension quiz, was associated with a lower model-based weight.^6^ Overall, the data show that in the absence of sufficient understanding, the standard analysis methods overestimate model-free influences on behaviour.

Model-free learning can explain many animal electrophysiology and human neuroimaging studies. A highly influential finding is that dopamine neurons respond to rewards consistently with the reward prediction errors obtained by temporal difference learning, a form of model-free learning.^37–39^ However, in the studies showing this response, there was no higher-order structure to the task—often, there was no instrumental task. Thus, model-based behaviour was indistinguishable from model-free. Conversely, when the task is more complicated, it has been found that the dopamine signal reflects knowledge of task structure (e.g.^40, 41^). A recent optogenetic study showed that dopamine signals are necessary and sufficient for model-based learning in rats,^42^ and consistent with this finding, neuroimaging studies in humans found that BOLD signals in striatal and prefrontal regions that receive strong dopaminergic innervation correlate with model-based prediction errors.^7, 43^ Moreover, although there is evidence that anatomically distinct striatal systems mediate goal-directed and habitual actions,^44^ to date there is no evidence for anatomically separate model-free and model-based systems.

Importantly, the two-stage task does not directly measure habits, and a lack of evidence for model-free learning does not imply humans don’t form or follow habits. Initial theoretical work proposed the model-based versus model-free distinction to formalize the distinction between goal-directed and habitual control.^4^ However, it is generally assumed that goal-directed actions can be based on model-free learning too. Similarly, there is an ongoing debate as to whether habits arise exclusively from model-free learning.^1, 4, 45–52^ In any case, two-stage task studies indicate that participants classified as primarily model-free are probably not acting on habits.

Apparently model-free participants behave inconsistently with the habitual tendencies that model-free learning supposedly indexes. A study by Konovalov and Krajbich^53^ combined eye-tracking with two-stage task choices to examine first-stage fixation patterns as a function of learning type. They divided participants into model-free and model-based, based on a median split of their estimated model-based weight (*w* = 0.3). They reported that when first-stage symbols were presented, model-based learners tended to look once at each symbol, as if they had already decided which to choose. In contrast, model-free learners tended to make more fixations, and their choices related more closely to fixation duration. These results suggest that model-free participants made goal-directed rather than habitual comparisons at the first stage. This is because similar patterns of head movements, presumably analogous to fixations, are seen when rats initially learn to navigate a maze.^54^ The head movements accompany hippocampal representations of reward in the direction the animal faces and are seen as evidence that the animals deliberate over choices in a goal-directed fashion. Humans also make more fixations per trial as difficulty increases in goal-directed paradigms.^55^ Notably, head movements and hippocampal place cell signaling cease once animals are extensively trained and act on habits.^54^

In contrast to habits, apparent model-free behaviour decreases with extensive training on the two-stage task. In general, the frequency and strength of habits increase with experience. However, Economides et al. showed that estimated model-free influence in human participants decreases over three days of training on the two-stage task.^56^ They also found that, after two days of training, participants remain primarily model-based even when performing a Stroop task in parallel. Both results raise questions about the effortfulness of model-based learning in the two-stage task. After all, its transition model may be tricky to explain, but it is easy to follow once understood. Rats also show primarily model-based behaviour after receiving extensive training on the two-stage task.^23^ Moreover, inactivation of the dorsal hippocampus or orbitofrontal cortex in the rats impaired model-based planning, but did not increase model-free influence, which remained negligible.^23^ Thus, while it is possible that humans and other animals may use model-free strategies in some cases, these results are difficult to reconcile with the idea of a competition between model-based and model-free systems for behavioural control.

Humans have been reported to arbitrate between model-based and model-free strategies based on both their accuracy and effort. Several lines of evidence indicate that humans and other animals generally seek to minimize physical and mental effort.^57^ Model-based learning is thought to require more effort than model-free learning, and a well-known aspect of the original two-stage task^7^ is that model-based learning does not lead to greater payoffs than model-free learning.^21, 29^ This has been hypothesized to lead participants to use a partially model-free strategy.^21, 29^ Previous studies tested this hypothesis by modifying the original two-stage task so that model-based strategies achieve more rewards.^21, 22, 24^ Participants appeared more model-based when a model-based strategy paid off more, and thus they concluded that participants will employ model-based learning if it is advantageous in a cost-benefit trade-off between effort and money. Our results and those from studies with extensive training^23, 58^ cannot be explained by such trade-offs. The magic carpet and spaceship tasks led to almost completely model-based behaviour, but had the same trade-offs as the original two-stage task.^7^ Similarly, the profitability of model-based learning does not change with experience. If anything, more experience should allow the agent to learn that the model-based strategy is no better than the model-free one if both are being computed in parallel. However, a possibility that merits further study is that giving causes for rare transitions reduced the subjective effort of forming the correct model.

Seemingly model-free behaviour may be reduced in all three sets of experiments through better understanding of the task. Obviously, improved instructions and more experience can lead to better understanding. In modified two-stage tasks that make model-based learning more profitable, the differential payoffs also provide clearer feedback to participants about the correctness of their models. Conversely, if both correct and incorrect models lead to the same average payoffs, participants may be slow to realize their mistakes. Of course, increased understanding and changes in cost-benefit ratios may jointly drive the increases in (correct) model-based behaviour, and additional data are needed to tease these possibilities apart.

It is not clear what is being measured by the two-stage task. Studies showed that IQ, working memory, processing speed, and dlPFC function are associated with model-based weights in the standard two-stage task with common instructions.^10–12, 59^ Therefore, determining and employing the correct model-based strategy probably relies on working memory and other PFC-mediated cognitive functions. Apparent model-free behaviour has been reported to correlate with compulsive symptoms.^6, 14, 60^ Given our current results, however, the conclusion that model-free learning and compulsive behaviours are linked in healthy populations or those with obsessive compulsive disorder should be drawn with caution (see also Supplementary Discussion).

Reward learning is one of the most widely studied processes in the neural and behavioural sciences. Researchers must use tools that can ascertain their mechanistic properties, their potential neural substrates, and how their dysfunction might lead to sub-optimal behaviour or psychopathology. The specification of algorithms for model-free versus model-based learning has advanced the study of reward learning. However, as Nathaniel Daw recently noted, “such clean dichotomies are bound to be oversimplified. In formalizing them, the [model-based]-versus-[model-free] distinction has also offered a firmer foundation for what will ultimately be, in a way, its own undoing: getting beyond the binary”.^28^ We believe that our current results are another strong indication that the time to move beyond oversimplified binary frameworks has come.

## Methods

### The magic carpet and spaceship tasks

24 healthy participants participated in the magic carpet experiment and 21 in the spaceship experiment. In both cases, participants were recruited from the University of Zurich’s Registration Center for Study Participants. The inclusion criterion was speaking English, and no participants were excluded from the analysis. No statistical methods were used to pre-determine sample sizes, but our sample sizes were based on our previous pilot studies using the two-stage task.^61^ The experiment was conducted in accordance with the Zurich Cantonal Ethics Commission’s norms for conducting research with human participants, and all participants gave written informed consent.

Participants first read the instructions for the practice trials and completed a short quiz on these instructions. For the magic carpet and spaceship tasks, 50 and 20 practice trials were performed respectively. Next, participants read the instructions for the main task, which was then performed. For the magic carpet task, they performed 201 trials and for the spaceship task, 250 trials. This number of trials excludes slow trials. Trials were divided into three blocks of roughly equal length. For every rewarded trial in the magic carpet or spaceship task, participants were paid CHF 0.37 or CHF 0.29 respectively. The total payment was displayed on the screen and the participants were asked to fill in a short questionnaire. For the magic carpet task, the questionnaire contained the following questions:

1. For each first-stage symbol, “What was the meaning of the symbol below?”
2. “How difficult was the game?” with possible responses being “very easy,” “easy,” “average,” “difficult,” and “very difficult.”
3. “Please describe in detail the strategy you used to make your choices.”

For the spaceship task, participants were only asked about their strategy. The questionnaire data are available in our Github repository along with all the code and the remaining participant data.

#### Magic carpet task description

Our magic carpet version of the two-stage task was framed as follows. Participants were told that they would be playing the role of a musician living in a fantasy land. The musician played the flute for gold coins to an audience of genies, who lived inside magic lamps on Pink Mountain and Blue Mountain. Two genies lived on each mountain. Participants were told that the symbol written on each genie’s lamp (a Tibetan character, see Figure 1C) was the genie’s name in the local language. When the participants were on a mountain, they could pick up a lamp and rub it. If the genie was in the mood for music, he would come out of his lamp, listen to a song, and give the musician a gold coin. Each genie’s interest in music could change with time. The participants were told that the lamps on each mountain might switch sides between visits to a mountain, because every time they picked up a lamp to rub it, they might put it down later in a different place.

To go to the mountains, the participant chose one of two magic carpets (Figure 1C). They had purchased the carpets from a magician, who enchanted each of them to fly to a different mountain. The symbols (Tibetan characters) written on the carpets meant “Blue Mountain” and “Pink Mountain” in the local language. A carpet would generally fly to the mountain whose name was written on it, but on rare occasions a strong wind blowing from that mountain would make flying there too dangerous because the wind might blow the musician off the carpet. In this case, the carpet would be forced to land instead on the other mountain. The participants were also told that the carpets might switch sides from one trial to the next, because as they took their two carpets out of the cupboard, they might put them down and unroll them in different sides of the room. The participants first did 50 “tutorial flights,” during which they were told the meaning of each symbol on the carpets, i.e., they knew which transition was common and which was rare. Also, during the tutorial flights, the participants saw a transition screen (Figure 1D), which showed the carpet heading straight toward a mountain (common transition) or being blown by the wind in the direction of the other mountain (rare transition). During the task, however, they were told their magic carpets had been upgraded to be entirely self-driving. Rather than drive the carpet, the musician would instead take a nap aboard it and would only wake up when the carpet arrived on a mountain. During this period a black screen was displayed. Thus, participants would have to figure out the meaning of each symbol on the carpets for themselves. The screens and the time intervals were designed to match the original abstract task,^7^ except for the black “nap” screen displayed during the transition, which added one extra second to every trial.

#### Spaceship task description

We also designed a second task, which we call the spaceship task, that differed from the original task reported in Daw et al.^7^ in terms of how the first stage options were represented. Specifically, there were four configurations for the first-stage screen rather than two. These configurations were represented as four different flight announcements displayed on a spaceport information board (Figure 4A). The task design was based on the common assumption that model-free learning is unable to generalize between distinct state representations.^15, 16, 21^ It has also been argued that reversals of the transition matrix should increase the efficacy of model-based compared to model-free learning.^29^ This type of reversal could happen between each trial in the spaceship task depending on which flight was announced. Thus, there are two reasons to expect that participants completing the spaceship task may be more model-based compared to the magic carpet task.

The spaceship task instructions stated that the participant would play the role of a space explorer searching for crystals on alien planets. These crystals possessed healing power and could be later sold in the intergalactic market for profit. The crystals could be found inside obelisks that were left on the surfaces of planets Red and Black by an ancient alien civilization. The obelisks grew crystals like oysters grow pearls, and the crystals grew at different speeds depending on the radiation levels at the obelisk’s location. There were two obelisks on each planet, the left and the right obelisk, and they did not switch sides from trial to trial. To go to planet Red or Black, the participant would use the left or the right arrow key to buy a ticket on a ticket machine that would reserve them a seat on spaceship X or Y. The buttons to buy tickets on spaceships X and Y were always the same. A board above the ticket machine announced the next flight, for example, “Spaceship Y will fly to planet Black.” Participants were told that the two spaceships were always scheduled to fly to different planets, that is, if spaceship Y was scheduled to fly to planet Black, that meant spaceship X was scheduled to fly to planet Red. Thus, if the announcement board displayed “Spaceship Y” and “Planet Black,” but they wanted to go to planet Red, they should book a seat on spaceship X.

After buying the ticket, the participant observed the spaceship flying to its destination. The participant was able to see that the spaceship would usually reach the planet it was scheduled to fly to, but in about one flight out of three the spaceship’s path to the target planet would be blocked by an asteroid cloud that appeared unpredictably, and the spaceship would be forced to do an emergency landing on the other planet. The precise transition probabilities were 0.7 for the common transition and 0.3 for the rare transition (Figure 1B). This transition screen was displayed during both the practice trials and the task trials (Figure 1D), and it explained to the participants why the spaceship would commonly land on the intended planet but in rare cases land on the other instead.

Thus, other than the four flight announcements, the spaceship task differed from the original two-stage task in that (1) the first-stage choices were labelled X and Y and were always displayed on the same sides, (2) for each choice, the participants were told which transition was common and which was rare, as well as the transition probabilities, (3) the participants saw a transition screen that showed if a trial’s transition was common or rare, (4) the second-stage options were identified by their fixed position (left or right), and (5) the time intervals for display of each screen were different. Many of these changes should facilitate model-based learning by making the task easier to understand.

### Simulations of model-based agents

The model-based agents described in the Results section were simulated and their decisions analysed by reinforcement learning model fitting. 1 000 agents of each type (original hybrid, unlucky symbol, and transition-dependent learning rates) performed a two-stage task with 1 000 trials, and the raw data were used to plot the stay probabilities depending on the previous trial’s outcome and reward. The hybrid reinforcement learning model proposed by Daw et al.^7^ was fitted to the data from each agent by maximum likelihood estimation. To this end, the optimization algorithm LBFGS, available in the PyStan library,^62^ was run 10 times with random seeds and for 5000 iterations to obtain the best set of model parameters for each agent. The three types of model-based agents we simulated are described below.

#### The hybrid algorithm

Daw et al.^7^ proposed a hybrid reinforcement learning model, combining the model-free SARSA(*λ*) algorithm with model-based learning, to analyse the results of the two-stage task.

Initially, at time *t* = 1, the model-free values 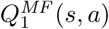 of each action *a* that can be performed at each state *s* are set to zero. At the end of each trial *t*, the model-free values of the chosen actions are updated. For the chosen second-stage action *a*_2_ performed at second-stage state *s*_2_ (the pink or blue states in Fig. 1A), the model-free value is updated depending on the reward prediction error, defined as 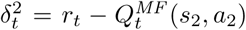, the difference between the chosen action’s current value and the received reward *r*_*t*_. The update is performed as

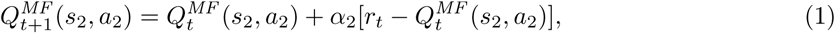

where *α*_2_ is the second-stage learning rate (0 *≤ α*_2_ *≤* 1). For the chosen first-stage action *a*_1_ performed at the first-stage state *s*_1_, the value is updated depending on the reward prediction error at the first and second stages, as follows:

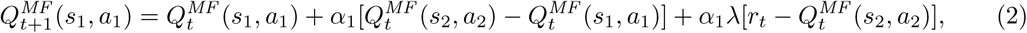

where *α*_1_ is the first-stage learning rate (0 *≤ α*_1_ *≤* 1), 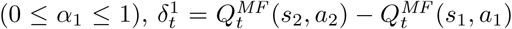 is the reward prediction error at the first stage, and *λ* is the so-called eligibility parameter (0 *≤ λ ≤* 1), which modulates the effect of the second-stage reward prediction error on the values of first-stage actions.

The model-based value 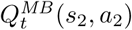 of each action *a*_2_ performed at second-stage state *s*_2_ is the same as the corresponding model-free value, i.e., 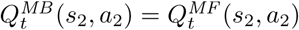. The model-based value of each first-stage action *a*_1_ is calculated at the time of decision making from the values of second-stage actions as follows:

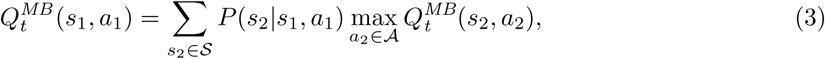

where *P* (*s*_2_|*s*_1_, *a*_1_) is the probability of transitioning to second-stage state *s*_2_ by performing action *a*_1_ at first-stage state *s*_1_, *S* = *{pink, blue}* is the set of second-stage states, and *A* is the set containing the actions available at that state.

The agent makes first-stage choices based on both the model-free and the model-based state-action pairs, weighted by a model-based weight *w* (0 *≤ w ≤* 1), according to a soft-max distribution:

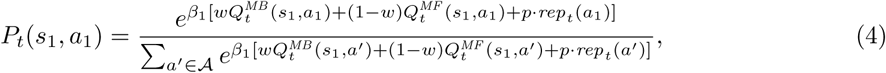

where *β*_1_ is the first-stage’s inverse temperature parameter, which determines the exploration-exploitation trade-off at this stage, *p* is a perseveration parameter that models a propensity for repeating the previous trial’s first-stage action in the next trial, and *rep*_*t*_(*a*′) = 1 if the agent performed the first-stage action *a*′ in the previous trial, and zero otherwise. Kool et al.^21^ have added an additional parameter to the hybrid model—the response stickiness *ρ*—and the above equation becomes

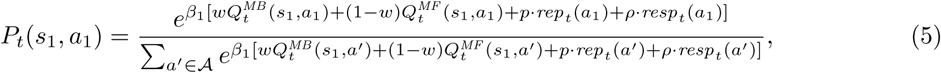

where the variable *resp*_*t*_(*a*′) is equal to 1 if *a*′ is the first-stage action performed by pressing the same key as in the previous trial, and zero otherwise.

Choices at the second stage are simpler, as the model-free and model-based values of second-stage actions are the same and there is no assumed tendency to repeat the previous action or key press. Second-stage choice probabilities are given as follows:

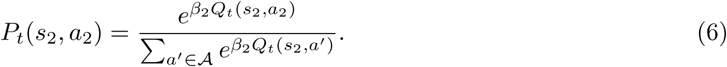

We propose two alternative algorithms below to demonstrate that model-based agents may be mistakenly classified as hybrid agents. These algorithms are based on the algorithm by Daw et al.^7^ detailed above, except that the inverse temperature parameter is the same for both stages (for simplicity because these models are only intended as demonstrations), the perseveration parameter *p* is equal to 0 (again, for simplicity), and the model-based weight *w* is equal to 1, indicating a purely model-based strategy.

#### The unlucky-symbol algorithm

We simulated an agent that believes a certain first-stage symbol is unlucky and lowers the values of second-stage actions by 50%. We reasoned that it is possible that an agent may believe that a certain symbol is lucky or unlucky after experiencing by chance a winning or losing streak after repeatedly choosing that symbol. Thus, when they plan their choices, they will take into account not only the transition probabilities associated to each symbol but also how they believe the symbol affects the reward probabilities of second-stage choices.

This model-based algorithm has three parameters: 0 *≤ α ≤* 1, the learning rate, *β >* 0, an inverse temperature parameter for both stages (for simplicity), and 0 *< η <* 1, a reduction of second-stage action values caused by choosing the unlucky symbol. The value of each first-stage action *a*_1_ is calculated from the values of second-stage actions as follows:

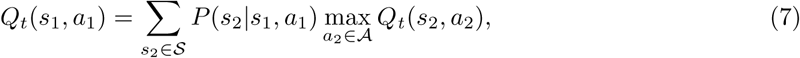

The probability of choosing a first-stage action is given by:

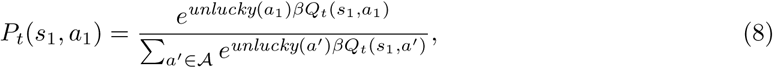

where *unlucky*(*a*) = *η* if the agent thinks action *a* is unlucky and *unlucky*(*a*) = 1 otherwise. Second-stage value updates and second-stage choices are made as described above for the original hybrid model. The probability of choosing a second-stage action is given by

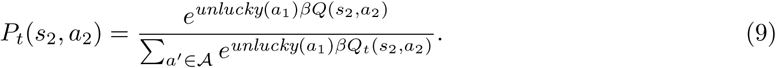

Learning of second-stage action values occurs as in the original hybrid model.

#### The transition-dependent learning rates (TDLR) algorithm

This is a simple model-based learning algorithm that has a higher learning rate after a common transition and a lower learning rate after a rare transition; hence, the learning rates are transition-dependent. The TDLR algorithm was inspired by debriefing comments from participants in a pilot study, which suggested that they assign greater importance to outcomes observed after common (i.e. expected) relative to rare transitions.

The TDLR algorithm has three parameters: *α*_*c*_, the higher learning rate for outcomes observed after common transitions (0 *≤ α*_*c*_ *≤* 1), *α*_*r*_, the lower learning rate for outcomes observed after rare transitions (0 *≤ α*_*r*_ *< α*_*c*_), and *β >* 0, an inverse temperature parameter that determines the exploration-exploitation trade-off. In each trial *t*, based on the trial’s observed outcome (*r*_*t*_ = 1 if the trial was rewarded, *r*_*t*_ = 0 otherwise), the algorithm updates the estimated value *Q*_*t*_(*s*_2_, *a*_2_) of the chosen second-stage action *a*_2_ performed at second-stage state *s*_2_ (pink or blue). This update occurs according to the following equation:

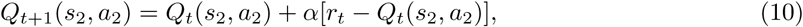

where *α* = *α*_*c*_ if the transition was common and *α* = *α*_*r*_ if the transition was rare. The value of each first-stage action *a*_1_ is calculated from the values of second-stage actions as follows:

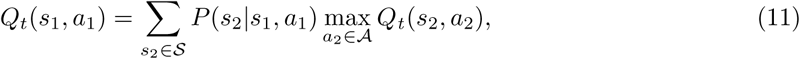

where *P* (*s*_2_|*s*_1_, *a*_1_) is the probability of transitioning to second-stage state *s*_2_ by performing action *a*_1_ at first-stage *s*_1_, *S* is the set of second-stage states, and *A* is the set of all second-stage actions. Choices made at first- or second-stage states are probabilistic with a soft-max distribution:

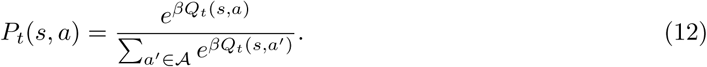

#### Simulation parameters

We simulated 1 000 purely model-based agents performing the two-stage task using each of the algorithms described above: (1) the original hybrid algorithm using a model-based weight *w* = 1 and *α*_1_ = *α*_2_ = 0.5, (2) the unlucky-symbol algorithm with *α* = 0.5 and *η* = 0.5, and (3) the TDLR algorithm with *α*_*c*_ = 0.8 and *α*_*r*_ = 0.2. For all agents, the *β* parameters had a value of 5.

#### The hybrid algorithm with a mistake probability

We developed a hybrid model with a mistake probability to analyse the symbol location effects in the common instructions data set. An additional parameter was added to the original hybrid reinforcement learning model: *ρ*^*i*^, the probability of making a mistake and making the wrong choice when the first-stage symbols switched sides from one trial to the next. Precisely, in trials with the first-stage symbols on different sides compared with the previous trials, let 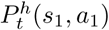 be the probability, according to the standard hybrid model, that the participant selected action *a*_1_ at the first-stage *s*_*i*_ in trial *t*. The same probability according the hybrid model with a mistake probability was given by

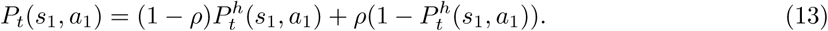

This model also assumes that the participant realized their mistake after making one and that action values were updated correctly.

### Analysis of the common instructions data

In,^21^ 206 participants recruited via Amazon Mechanical Turk performed the two-stage task for 125 trials. The behavioural data were downloaded from the first author’s Github repository (https://github.com/wkool/tradeoffs) and reanalysed by logistic regression and reinforcement learning model fitting, as described below.

### Logistic regression of consecutive trials

This analysis was applied to all behavioural data sets. Consecutive trial pairs were analysed together, or first divided into subsets, depending on the presentation of first-stage stimuli, then analysed separately. The analysis employed a hierarchical logistic regression model whose parameters were estimated through Bayesian computational methods. The predicted variable was *p*_stay_, the stay probability, and the predictors were *x*_*r*_, which indicated whether a reward was received or not in the previous trial (+1 if the previous trial was rewarded, −1 otherwise), *x*_*t*_, which indicated whether the transition in the previous trial was common or rare (+1 if it was common, −1 if it was rare), the interaction between the two. Thus, for each condition, an intercept *β*_0_ for each participant and three fixed coefficients were determined, as shown in the following equation:

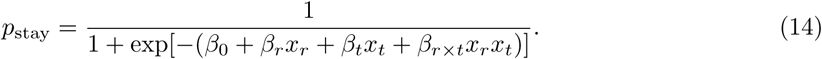

For each trial pair, the variable *y* was equal to 1 if the agent chose in the next trial the same first-stage action as in the previous trial (a “stay” choice) or equal to 0 if the agent chose a different first-stage action (a “switch” choice). The distribution of *y* was Bernoulli(*p*_stay_). The distribution of the 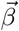 vectors was 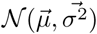. The hyperparameters 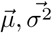 were given vague prior distributions based on preliminary analyses—the 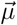 vectors’ components were given a *N* (*µ* = 0, *σ*^2^ = 25) prior, and the 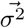 vector’s components were given a Cauchy(0, 1) prior. Other vague prior distributions for the model parameters were tested and the results did not change significantly.

To obtain parameter estimates from the model’s posterior distribution, we coded the model into the Stan modeling language^63, 64^ and used the PyStan Python package^62^ to obtain 60 000 samples of the joint posterior distribution from four chains of length 30 000 (warmup 15 000). Convergence of the chains was indicated by 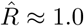 for all parameters.

### Fitting of reinforcement learning models

The hybrid and correct model-based reinforcement learning model proposed by Daw et al.^7^ were fitted to all data sets (common instructions, magic carpet, and spaceship). To that end, we used a Bayesian hierarchical model, which allowed us to pool data from all participants to improve individual parameter estimates. For the analysis of the spaceship data, four distinct first-stage states were assumed, corresponding to the four possible flight announcements (Figure 4A).

The parameters fitted for the standard hybrid and correct model-based algorithms were as follows. The parameters of the hybrid model for the *i*th participant were 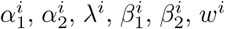, and *ρ*^*i*^. Vectors

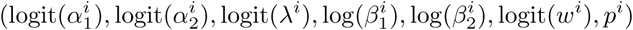

were given a multivariate normal distribution with mean ***µ*** and covariance matrix **Σ** and obtained for each participant. These transformations of the parameters were used because the original values were constrained to an interval and the transformed ones were not, which the normal distribution requires. The correct model-based algorithm had the same parameters except *α*_1_, *λ*, and *w*. The model’s hyperparameters were given weakly informative prior distributions. Each component of ***µ*** was given a normal prior distribution with mean 0 and variance 5, and **Σ** was decomposed into a diagonal matrix ***τ***, whose diagonal components were given a Cauchy prior distribution with mean 0 and variance 1, and a correlation matrix **Ω**, which was given an LKJ prior^65^ with shape *ν* = 2.^64^ This model was coded in the Stan modelling language^63, 64^ and fitted to each data set using the PyStan interface^62^ to obtain a chain of 40 000 iterations (warmup: 20 000) for the common instructions data set and 80 000 iterations (warmup: 40 000) for the magic carpet and spaceship data sets. Convergence was indicated by 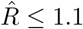 for all parameters.

The same procedure above was performed to fit a hybrid model with a mistake probability to the common instructions and magic carpet data sets. For that model, the data from each participant were described by a vector,

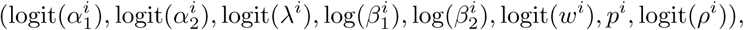

where the additional parameter, logit(*ρ*^*i*^), is the mistake probability after a side switch.

The hybrid and model-based algorithms were also fitted to data by maximum likelihood. They were first coded in the Stan modelling language^63, 64^ and fitted 1000 times (for robustness) to each participant’s choices using LBFGS algorithm implemented by Stan through the PyStan interface.^62^

### Model comparisons

To calculate PSIS-LOO scores for model comparison, each model was first fitted simultaneously to all behavioural data in each condition, as described above. Then, the log-likelihood of every trial was obtained for each iteration and used to calculate the PSIS-LOO score (an approximation of leave-one-out cross-validation) of each model (considering all trials from all participants together) or of each model for each participant (considering only the trials from that participant). To this end, the loo and compare functions of the loo R package were employed.^66^

To calculate AIC scores, we fit the hybrid and the model-based reinforcement learning models to all data sets by maximum likelihood estimation for each participant, as described above. The logistic regression model was fitted to the data from each participant using the statsmodels library.^67^ AIC scores were then calculated from the log-likelihood of the model fit for each participant and summed across each data set and model.

For the comparisons between the correct model-based, hybrid, and logistic regression models, the AIC and PSIS-LOO scores were computed for the first-stage choices only. This is because the logistic regression model is only fit to first-stage choices, and the model-free and model-based algorithms only generate different behaviour at the first stage, but not at the second.

## Data availability statement

The data obtained from human participants are available at https://github.com/carolfs/muddled_models

## Code availability statement

All the code used to perform the simulations, run the magic carpet and the spaceship tasks, and analyse the results are available at https://github.com/carolfs/muddled_models

## Acknowledgements

We would like to thank Giuseppe M. Parente for the wonderful illustrations used in the spaceship and magic carpet tasks, Susanna Gobbi, Gaia Lombardi, and Micah Edelson for many helpful discussions and ideas, the participants at the NYU Neuroeconomics Colloquium for useful feedback, and Peter Dayan, Wouter Kool, Arkady Konovalov, Ian Krajbich, and Stephan Nebe for helpful comments on early drafts of this manuscript. Note that our acknowledgment of their feedback does not imply that these individuals fully agree with our conclusions or opinions in this paper. We would also like to acknowledge Wouter Kool, Fiery Cushman, and Sam Gershman for making the data from their 2016 paper openly available at https://github.com/wkool/tradeoffs. This work was supported by the CAPES Foundation (https://www.capes.gov.br, grant number 88881.119317/2016-01) and the European Union’s Seventh Framework programme for research, technological development and demonstration under grant agreement no 607310 (Nudge-it). The funders had no role in study design, data collection and analysis, decision to publish or preparation of the manuscript.

## Author contributions

C.F.S. and T.A.H. designed the tasks and novel computational models. C.F.S. programmed the tasks, collected the data, and performed the analyses with input from T.A.H. C.F.S. and T.A.H. wrote the manuscript.

## Competing interests

The authors declare no competing interests.

## Notes

https://github.com/carolfs/muddled_models

