## Supplementary Material for "Humans are primarily model-based learners in the two-stage task"

#### Supplementary Methods

##### Further details on the instructions used in the common instructions experiment

Kool et al.[1] also framed the two-stage task within the context of a story. However, this story omitted explanations for some of the task events covered by our instructions. These omissions may be responsible for some of the differences in behaviour we observe across the two studies. Following Decker et al.,[2] they explained to participants that the first stage consisted of choosing a spaceship to travel to one of two planets, where they would meet a pair of friendly aliens, who might or might not share with them some space treasure from a mine. The common instructions stated that "on each trial, you will first choose which spaceship to take. The spaceships can fly to either planet, but one will mostly fly to the green planet, and the other mostly to the yellow planet. The planet a spaceship goes to most won't change during the game. Pick the one that you think will take you to the alien with the best mine, but remember sometimes you'll go to the other planet!" Participants were not told why a spaceship frequently flies to one planet, but sometimes flies to the other. In our daily lives, when we take a bus, plane, train to a certain location, we expect to eventually arrive at that location, and if there is a deviation from the intended destination, we expect there to be a good and clearly communicated reason for this change. The fact that no reasons are given for the spaceships' occasional deviations from their normal paths may have led participants to begin hypothesizing on their own about potential causes for this behaviour. It is plausible that some participants might have imagined that the reward outcomes were somehow linked to transition types even though no such relationship exists.

Participants were also not told why the spaceships and aliens might be on different sides of the screen in different trials. Here again, they might have created their own explanations to fill in these gaps. One set of potential self-generated explanations might cause participants to form an incorrect model of the task in which the side of the screen the spaceships appear on influences the rewards. If participants behaved in accordance with this incorrect model, then the standard analyses of two-stage task choices will incorrectly indicate that there was a significant model-free component to their choices. However, in reality participants may just have formed an inaccurate mental model of the task.

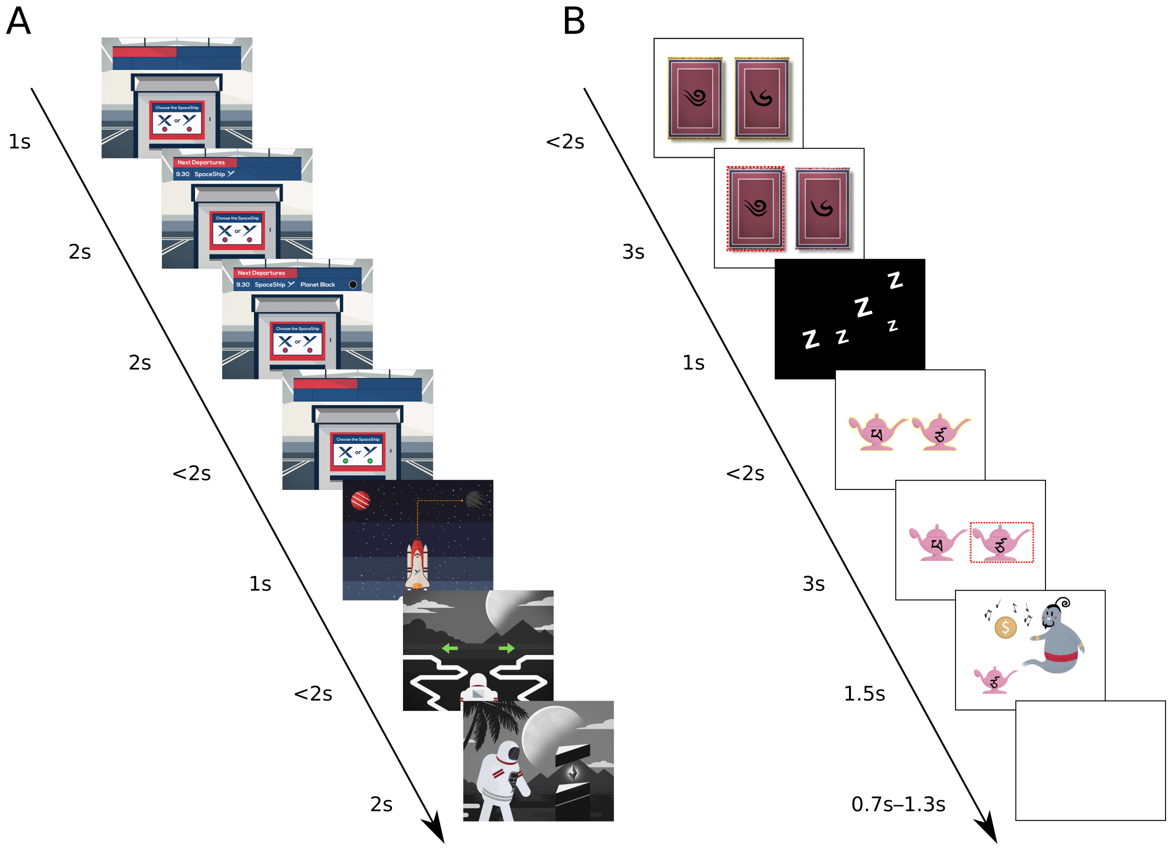


Figure 7: Timelines of the spaceship and magic carpet task. Each box depicts an event within the spaceship or magic carpet tasks. The duration of each even is given in seconds on the left. A) In the spaceship task, the 1st screen simply indicates that a new trial has begun. The 2nd and 3rd screens represent the initial state. At the 4th screen, the participant has up to 2 seconds to indicate her choice. The common or rare transition is shown on the 5th screen. The second-stage state was indicated by the background color (black, red) and the choice by the green left and right arrows on the 6th screen. The 7th and final screen in a trial revealed whether or not a reward was delivered. After feedback, the task advanced directly to the next trial. B) The magic carpet task was designed to closely mimic the original, abstract version of the two-stage task while still allowing for story-based instructions that included causes and effects for all task events. Thus, we used the same Tibetan characters from the original task, made them into labels for magic carpets and genies rather than simply identifying colored squares. In the magic carpet task, the 1st screen represented the initial state and first-stage choice. Participants had up to 2s to make this choice. On the second screen, the chosen option was highlighted for 3 seconds. Next, a "nap" screen was shown for 1s while the magic carpet automatically took the participant to one of the two mountains. Although participants saw the common or rare transition screens depicted in Figure 1D during the practice trials, the transitions were not shown during the main task to make it more comparable with previous versions. The second-stage state (blue, pink) and choice were indicated by the pink or blue lamps on the right and left side of the 4th screen. The participant had up to 2s to make her choice. The 5th screen highlighted the chosen lamp/genie for 3s. The 6th and final screen in a trial revealed whether or not a reward was delivered. After reward feedback, there was a blank screen for 0.7–1.3s before the next trial began.

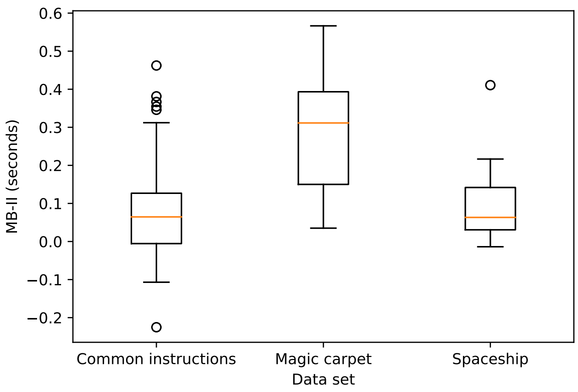


Figure 8: Distribution of MB-II scores in the common instructions, magic carpet, and spaceship data sets. MB-II scores are defined as the mean reaction time at the second stage after a rare transition minus the mean reaction time at the second stage after a common transition.[3] They have been considered a "reinforcement-learning-model-agnostic estimate" of model-based behaviour, under the assumption that more model-based agents choose faster after a common transition and slower after a rare transition compared to more model-free agents. Each box displays the 0.05, 0.25, 0.5, 0.75, and 0.95 quantiles of each distribution. The circles represent outliers, defined as participants whose MB-II scores fall outside the 0.05–0.95 quantile range. MB-II scores for the magic carpet task are higher compared to the common instructions task. However, MB-II scores for the spaceship task are lower compared to the magic carpet task. This is likely because in the spaceship task, after participants make their choice at the first stage but before they arrive at the second stage, they see a screen that presents information about the transition and second-stage state where they are about to arrive (Figure 7).

### Supplementary Results

#### Reaction times for second-stage choices as a function of transition type

Previous studies have shown that human participants' reaction times at the second stage correlate with other measures of model-based behaviour.[2,3] Participants who appear more model-based are slower at making second-stage choices after a rare transition and faster after a common transition, compared to participants who appear more model-free. We analysed second-stage reaction time data from the common instructions, magic carpet, and spaceship data sets to check for the effect of transition. In all of our other analyses, participants in the magic carpet and spaceship tasks, compared to the common instructions task, displayed higher measures of correct model-based behaviour. We would thus expect this difference to be consistent with second-stage reaction times after common versus rare transitions. We calculated each participant's MB-II score, defined as the mean reaction time after a rare transition minus the mean reaction time after a common transition.[3] We observed that indeed participants in the magic carpet task obtained higher MB-II scores compared to the common instructions task (Figure 8). The MB-II scores of participants in the spaceship task were not as high as the magic carpet, likely because they received information about the transition and second-stage state for one second before arriving at the second stage, as shown in Figure 7.

We note, however, a caveat about using reaction times as evidence of "model-basedness" or understanding of the task structure: a strong effect of transitions on reaction times only indicates that participants understood the transition structure, not that they have the correct model of task or specifically how their models are incorrect. Recall that our simulated example models both include the correct transition structure, but still produce behavioural patterns that could be mistaken for hybrid.

#### Transition main effects

The data from the magic carpet and spaceship tasks show small, but significant effects of transition on stay probabilities (Figure 3B). This coefficient indicates that the probability of repeating the same first-stage action increases after experiencing a common transition to the second stage. Transition effects have not received much attention in the previous literature on two-stage tasks. However, they

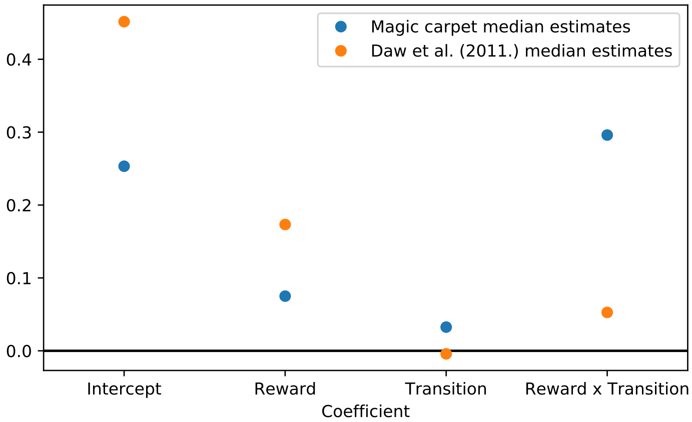


Figure 9: Some hybrid agents display a main effect of transition in a logistic regression analysis. The logistic regression coefficients in blue were generated by a hybrid agent using the median parameter values from the magic carpet task, which are $\alpha_1 = 0.67, \alpha_2 = 0.91, \lambda = 0.6, \beta_1 = 6, \beta_2 = 2.73, w = 0.79, p = 0.03$. For comparison, the coefficients in orange were generated by a hybrid agent using the median parameter values from the original paper by Daw et al.,[4] which are $\beta_1 = 5.19, \beta_2 = 3.69, \alpha_1 = 0.54, \alpha_2 = 0.42, \lambda = 0.57, p = 0.11, w = 0.39$. All simulated agents performed the two-stage task for 201 trials.

are consistent with either model-free or model-based behaviour under certain combinations of the learning and choice parameters in those algorithms.

Note that the prediction in Figure 4B, which shows a main effect of transition, was generated by the same hybrid model proposed by Daw et al.[4] While this hybrid model predicts a null transition effect for some parameter combinations, this is not universally true for all values of the learning rate ($\alpha$), eligibility trace ($\lambda$), softmax temperature ($\beta$), and model based weight ($w$) parameters. Figure 9, for instance, shows in blue the logistic regression coefficients generated by a hybrid agent using the median parameter values from our magic carpet task. For comparison, the fits for an agent using the median values from the original paper by Daw et al.[4] are shown in orange. In summary, both the model-free, hybrid and correct model-based algorithms can each generate both zero and non-zero transition effects in a logistic regression analysis of choice behaviour.

#### Even with the new instructions, the original hybrid model still does not capture all relevant features of human participants' behaviour

Despite the fact that our more comprehensive, story-based instructions seemed to alleviate many misconceptions of the two-stage task, participants' behaviour still did not strictly follow the assumptions inherent in the standard logistic regression or hybrid reinforcement learning model analyses. These deviations were readily apparent when we analysed the data from the magic carpet and spaceship task using a modified "many-trials-back" logistic regression model of the type first proposed by Miller et al.[5] Those authors showed that adding reward and transition predictors from many previous trials is a more robust method for analyzing behaviour on two-stage tasks. Here, we used a similar analysis that is a linear transformation of their method and thus includes and utilizes the same information (see Figure 10 and the next section for details).

We used this many-trials-back regression method in order to try and further understand our participants' strategies during the two-stage task. First, we fit the many-trials-back logistic regression directly to the empirical data from the magic carpet and spaceship tasks. Second, we fit the standard hybrid model[4] to the magic carpet and spaceship task data and then generated simulated choices for every participant from the standard hybrid model using the participants' maximum likelihood parameters. Ideally, when the hybrid learning model is run with these fitted parameters, it should make decisions in a way that resembles the respective participant's decisions. How well the simulated and empirically observed patterns of choices match up is an indication of the extent to which the hybrid model is able to explain and reproduce participants' behaviour.[6] The results of this procedure are

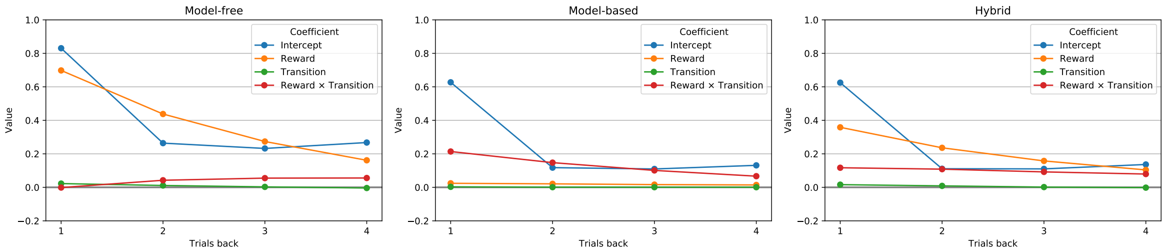


Figure 10: These three plots show the results from a modified "many-trials-back" logistic regression analysis of simulated reinforcement learning agents. The many-trials-back logistic regression we compute here is based on the analysis proposed by Miller et al.[7] The three plots show that our modified formulation of the many-trials-back regression can distinguish between model-free, model-based, and hybrid reinforcement learning agents. We simulated a model-free agent, a model-based agent, and a hybrid agent performing the two-stage task with standard hybrid model proposed in.[4] The model-based weights for each agent were 0, 1, and 0.5, respectively. After simulating each agents' behaviour, we analysed the resulting choices using the many-trials-back logistic regression. The model-free agent is characterized by a positive, decreasing reward effect. The model-based agent is characterized by a positive, decreasing reward by transition interaction effect. The hybrid agent exhibits both effects, as expected. Hence, our modification of Miller et al.'s method is, indeed, able to differentiate between these types of agents when model-based and hybrid agents use the correct model of the task. Error bars are not shown because they are very small due to the large number of data points.

shown in Figure 11.

There were two main differences between the results from empirical data and the standard hybrid model simulations: (1) the empirical reward by transition interaction is more than twice as large than the effect predicted by the hybrid model for the first two trials back, and (2) the hybrid model predicts a near zero effect of transition for all trials, but the empirical results exhibit a negative transition effect for the second and third trials back. These differences suggest that participants use a model of the task that differs substantially from the standard hybrid model. In particular, the negative transition effect suggests that participants may have employed a directed exploration strategy at the first stage of the two-stage decision task. This is because a negative transition effect for a past trial means participants are more likely to switch their first-stage choices after experiencing common than rare transitions. This pattern of behaviour suggests that participants actively avoided going to the same second-stage state several times in a row. In fact, many participants wrote in the debriefing questionnaire that they tried to go to a second-stage state two or three times in a row, and then intentionally aimed for the other state. The standard hybrid model cannot account for this type of behaviour.

#### Logistic regression analyses using many trials back

Miller et al.[5] have proposed a "many-trials-back" logistic regression analysis for quantifying behaviour on two-stage tasks and have shown it to be effective and, in fact, more robust than standard techniques in distinguishing between different variants of model-free, model-based, and hybrid behaviour in simulated agents. Here, we use an analogous method to determine if and how participants' deviate from the correct task model due to misunderstanding the task, boredom, exploration, etc.

Our version of the many-trials-back analysis works as follows. We estimated the effect of reward and transition in the past $T = 4$ trials on the first-stage choice made in the next trial $t'$. The first-stage choice for all trials were coded as $+1$ if it was "right" for the spaceship task and a certain symbol (the same for all participants) for the magic carpet task and $-1$ otherwise; a reward in trial $t$ was coded as $r_t = +1$ if the trial was rewarded and $r_t = -1$ otherwise, and transitions were coded as $\tau_t = +1$ if the transition was common and $\tau_t = -1$ if it was rare. Let $y$ be the first-stage choice on trial $t'$ and, for each of the $T$ previous trials $t$, $x_t$ be the first-stage choice made in that trial. The logistic regression

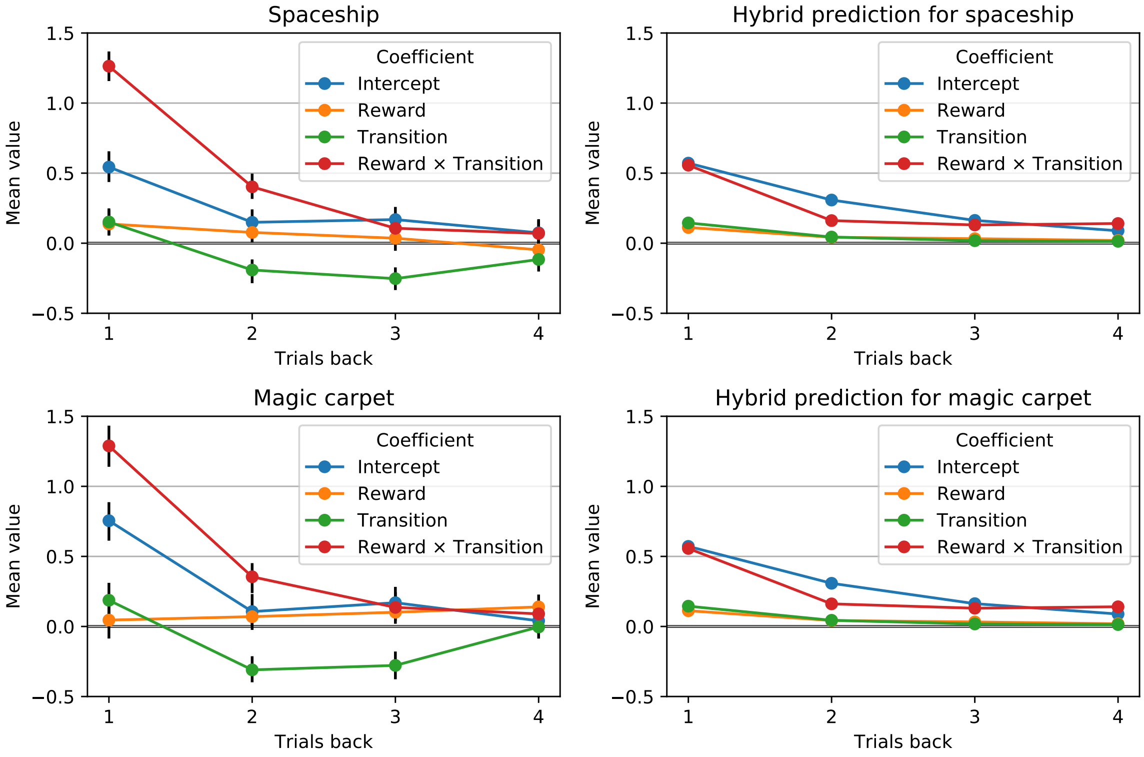


Figure 11: These four plots show the logistic regression results from the "many trials back" analysis. In this case many = 4 trials back. The plots in the left column show the experimental results from the magic carpet and spaceship tasks. The plots in the right column show the predicted results for the magic carpet and spaceship tasks, obtained by fitting the hybrid reinforcement learning model to the data, and then simulating agents using the fitted parameters to perform 1000 trials of each task. Error bars are not shown for the simulated results because they are very small due to the large number of data points. Error bars for the experimental results represent the 95% highest density intervals.

model is

$$\Pr(y=+1) = \frac{1}{1+\exp[-(\sum_{t=1}^{T}\beta_0^{t'-t}x_{t'-t}+\beta_x^{t'-t}r_{t'-t}x_{t'-t}+\beta_\tau^{t'-t}\tau_{t'-t}x_{t'-t}+\beta_{x\times\tau}^{t'-t}r_{t'-t}\tau_{t'-t}x_{t'-t})]}, \tag{1}$$

where $\Pr(y = +1)$ is the probability that the participant chose the choice encoded as +1 in the next trial, $\beta_0^{t'-t}$ is a coefficient that indicates the overall influence of the first-stage choice made in trial $t' - t$ on the first-stage choice made in trial $t'$, $\beta_r^{t'-t}$ is a coefficient that indicates the reward effect for trial $t' - t$, $\beta_\tau^{t'-t}$ is a coefficient that indicates the transition effect for trial $t' - t$, and $\beta_{r\times\tau}^{t'-t}$ is a coefficient that indicates the reward by transition interaction effect for trial $t' - t$.

A hierarchical Bayesian approach was used to perform the "many-trials-back" analysis on the magic carpet and spaceship data sets. To generate the hybrid model's predicted results for that analysis, the hybrid model was simulated using each participant's maximum likelihood parameters in 10 000 simulations. For each simulation, the logistic regression coefficients were obtained by fitting the hierarchical logistic regression model described above to the data and calculating the maximum a posteriori point of its posterior distribution, via PyStan's LBFGS algorithm with a random seed and 5000 iterations.[8]

### Supplementary Discussion

In the case of OCD, is the two-stage task picking up alterations in how distinct habitual (indexed by model-free learning) and goal-directed (model-based learning) systems interact to control behaviour? Or are differences in two-stage choices driven by the ability to comprehend the task structure, create and maintain accurate mental models of it, and use these models to make decisions? It is certainly possible that OCD patients and other individuals with sub-clinical compulsive symptoms do indeed use more model-free learning during the two-stage task. However, a plausible alternative explanation for the correlations with model-free indexes in the two-stage task is that compulsive individuals form and continue to follow inaccurate models of the world. Patients with OCD have been found to be impaired in causal reasoning between actions and outcomes,[9] deficits that would likely impair the ability to form accurate task models. In fact, individuals with OCD have been reported to be impaired in multiple measures of cognitive flexibility, but the fundamental reasons for these impairments remain unclear.[10,11] Our current findings do not change the fact that behaviour on the two-stage task is correlated with compulsive symptoms, but they do indicate a need to continue investigating the underlying cause for these correlations.
